# Multi- and transgenerational disruption of maternal behavior and female puberty by Endocrine Disrupting Chemical (EDC) mixture exposure

**DOI:** 10.1101/2020.06.26.172965

**Authors:** David López-Rodríguez, Carlos Francisco Aylwin, Virginia Delli, Elena Sevrin, Marzia Campanile, Marion Martin, Delphine Franssen, Arlette Gérard, Silvia Blacher, Ezio Tirelli, Agnès Noël, Alejandro Lomniczi, Anne-Simone Parent

## Abstract

Female reproductive development and maternal behavior are two intertwined phenotypes centrally controlled by the hypothalamus. Endocrine disrupting chemicals (EDC) can alter these processes especially when animals are exposed during development. We propose the concept that developmental exposure to a low environmentally relevant dose of EDC mixture induces a transgenerational alteration of female rat pubertal timing and ovarian physiology throughout epigenetic reprograming of hypothalamic *Kiss1, Esr1* and *Oxt1* loci. Such exposure also caused a multigenerational reduction of maternal behavior induced by the loss in hypothalamic dopaminergic signaling. Our results identify the hypothalamic Polycomb Group of epigenetic repressors as actors of this mechanism of transgenerational reproductive disruption. Using a cross-fostering approach, we identified that while the reduction in maternal phenotype was normalized in EDC exposed pups raised by unexposed dams, no reversal of the pubertal phenotype was achieved, suggesting a germline transmission of the reproductive phenotype.

## INTRODUCTION

Endocrine Disrupting Chemicals (EDC) impact populations as much as individuals given their environmental ubiquity^1^. Among 85,000 chemicals in use, 1000 have been identified as having the ability to disrupt normal endocrine function^2^. Developmental exposure to EDC has been associated with increased risk of genital malformations, hypofertility and testis cancer in human and rodent males, increased risk of breast cancer and metabolic syndrome in adulthood, as well as alterations of pubertal timing and ovarian function^3^. Fetal development as well as childhood and adolescence are known windows of sensitivity to chemical exposure, which can alter the developmental trajectory and lead to long-lasting dysfunction^1,4^. In addition to the direct effects of EDC on individuals, studies have revealed that early exposure affects the health of subsequent generations through epigenetic (transgenerational) mechanisms that alter the germ line methylome, affecting pubertal timing, ovarian follicle development, stress responsiveness and mate preference in subsequent generations^5,6^. Alternatively, a non-genetic transmission of parental traits has been shown to propagate multigenerationally^7^. Maternal care affects offspring development from early postnatal life, inducing somatic changes that lead to the propagation of the same behavioral phenotype such as increased stress responsiveness or low licking and grooming behavior^8^.

In mammals, sexual maturation is driven by increased pulsatile secretion of hypothalamic gonadotropin-releasing hormone (GnRH) released into the portal vasculature that feeds into the pituitary gland, ultimately increasing pulsatile release of luteinizing hormone (LH) from pituitary gonadotrophs into the peripheral circulation, inducing ovarian steroidogenesis and ovulation^9^. During the prepubertal period, the secretory activity of GnRH neurons is under predominant trans-synaptic inhibitory control provided by GABAergic, Opiatergic and RFamide inputs into the GnRH neuronal network^10^. At puberty this inhibition is lifted while a concomitant increase in excitatory inputs to the GnRH network is provided by Glutamatergic and Kisspeptidergic neurons^11,12^. Recent evidence suggests that this trans-synaptic regulatory mechanism is controlled by a molecular switch that regulates the timing of puberty. This epigenetic switch coordinates the transcriptional activity of arcuate nucleus (ARC) kisspeptin neurons implicated in stimulating GnRH release^13,14^. Before puberty, the transcriptional activity of *Kiss1*, the gene encoding for kisspeptins, is downregulated by members of the Polycomb Group (PcG) of transcriptional repressors, by catalyzing the trimethylation of histone 3 at lysine 27 (H3K27me3), a histone mark associated with gene silencing^14^. As puberty approaches, the PcG is evicted from the *Kiss1* promoter and the Trithorax Group (TrxG) of epigenetic activators is recruited, resulting in increased histone methylation at lysine 4 (H3K4me3) and acetylation at lysine 27 (H3K27ac) to the *Kiss1* promoter/enhancer regions respectively, leading to an increase in *Kiss1* mRNA transcription^13^.

Here we propose the concept that developmental exposure to an EDC mixture induces a transgenerational alteration of female pubertal timing and a multigenerational decrease in maternal behavior. In the current study, female rats (F0) were exposed to a mixture of 13 of the most prevalent EDC present in the human body at relevant exposure concentrations from 2 weeks before gestation until the end of lactation. The four subsequent generations were evaluated for sexual maturation and maternal behavior. Our data show that gestational and lactational exposure to an environmentally relevant EDC mixture alters maternal behavior in F1 trough F3 generations and transgenerationally affects sexual development by epigenetic reprogramming of the hypothalamus. While F2, F3 and F4 females display delayed puberty and abnormal estrous cycles, such changes are not detected in F1 *in vitro* and lactationally exposed animals. As the phenotype appears at the F2 generation, these data suggest that germline exposure is required to disrupt reproductive development. These phenotypes are associated with alterations in both transcriptional and histone posttranslational modifications of hypothalamic genes involved in dopamine signaling and GnRH neuron pulsatility control. By using a cross-fostering paradigm, we show that the reproductive alterations are transmitted through a germline-dependent mechanism, while the alteration in maternal behavior is, at least in part, produced by the direct exposure of the fetus to the EDC mixture.

## RESULTS

### Exposure to EDC mixture alters pubertal onset, estrous cycle and folliculogenesis throughout generations

Sexual maturation was followed for 4 generations (F1 to F4) of female rats after exposure of F0 females to a mixture of 13 anti-androgenic and estrogenic EDC or corn oil (vehicle) (Fig. 1). F0 females were exposed orally for 2 weeks before and throughout gestation until weaning. While F1 females, which were directly exposed to EDC *in utero* (F1-EDC), had normal pubertal timing (Fig. 2a) determined by age at vaginal opening; F2, F3 and F4 females had significantly delayed vaginal opening (Fig. 2b-d). Maturation of GnRH secretion preceding puberty is characterized by a reduction of GnRH interpulse interval between P15 and P25 in hypothalamic explants incubated individually^15^. While GnRH interpulse interval was not affected in F1-EDC at P20 (Fig. 2a), it was significantly increased in F3-EDC females (Fig. 2c), suggesting a delayed maturation of GnRH secretion, consistent with a delayed onset of puberty. In addition, estrous cyclicity was disrupted in F2 and F3-EDC females (Fig. 3b-c left) with a significant decrease in the proportion of females showing regular cycles, characterized by increased time spent in estrus and reduced time in diestrus. Additionally, F3-EDC females displayed a decreased time spent in proestrus. All these data show globally a sign of subfertility.

**Figure 1.**
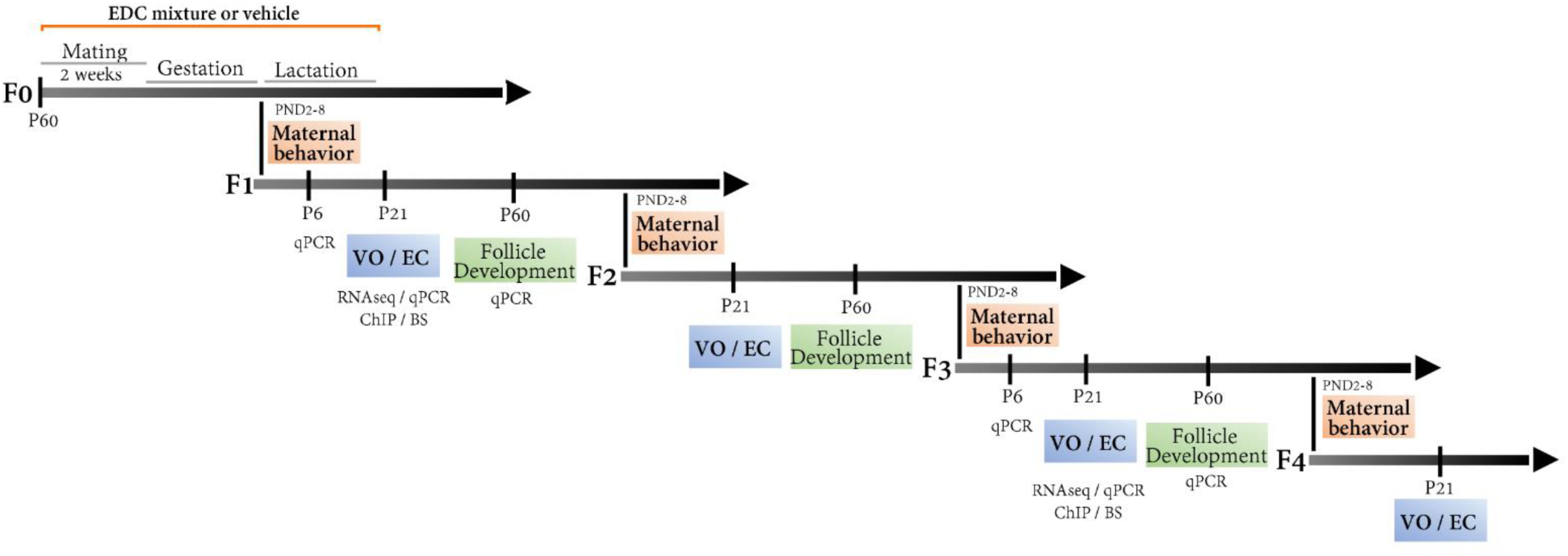
Multi- and transgenerational design of an EDC mixture exposure on maternal behavior and sexual maturation. Female rats (F0) were exposed to a mixture of 13 of the most prevalent EDC present in the human body at relevant exposure concentrations from 2 weeks before gestation until the end of lactation. The four subsequent generations were evaluated for sexual maturation (vaginal opening, GnRH interval interpulse, estrous cycle and folliculogenesis) and maternal behavior (from P2 to P8). Massive parallel RNA sequencing was carried out using mediobasal hypothalamic (MBH) explants from the F1 and F3 generation to decipher direct (F1) versus transgenerational (F3) target genes of the EDC mixture exposure, followed by qPCR validation at three time points, P6, 21 and 60. Target genes were studied for histone posttranslational modifications and DNA methylation using chromatin immunoprecipitation (ChIP) and bisulfite sequencing, respectively. V.O: vaginal opening, E.C: estrus cycle.

**Figure 2.**
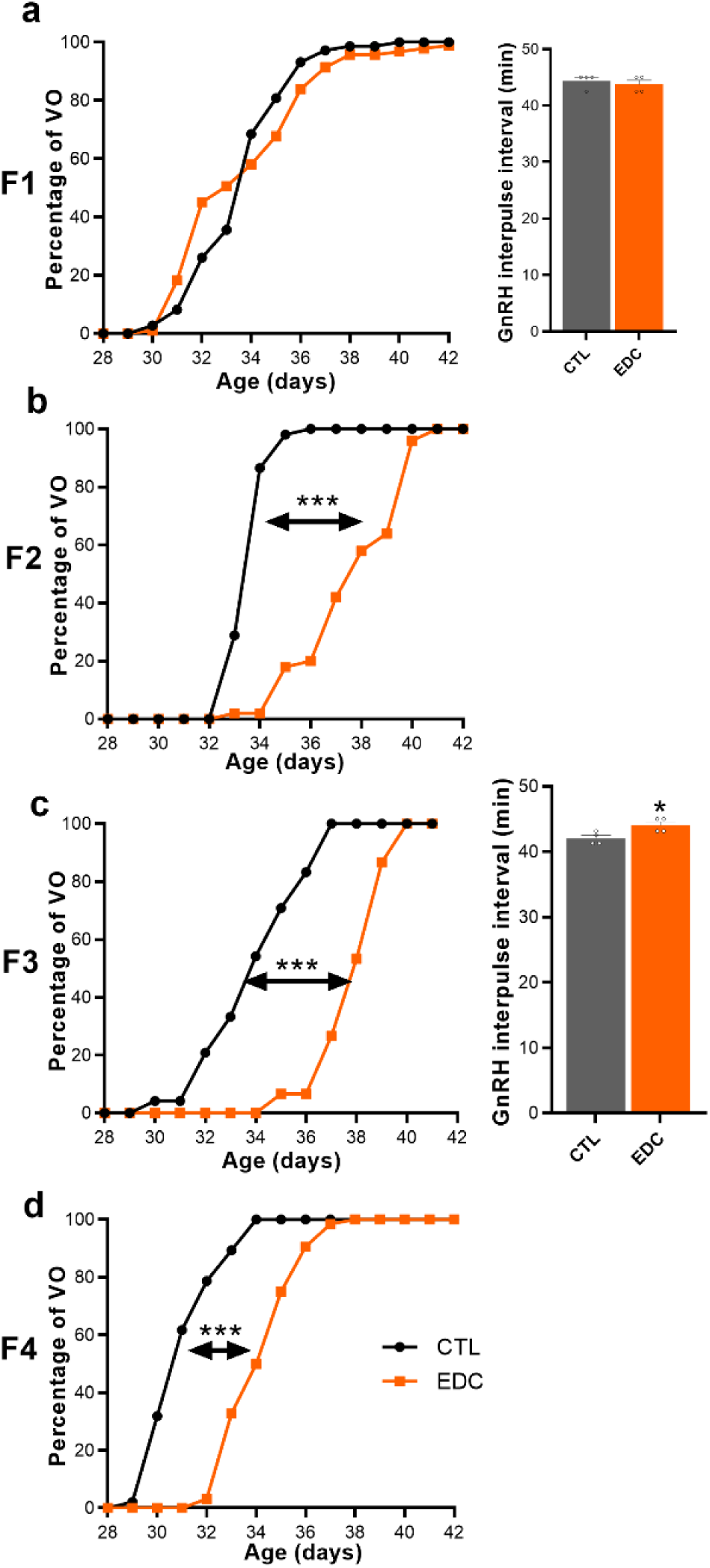
Pubertal timing (vaginal opening) and GnRH interpulse interval across generations (F1-F4 generation) after an EDC mixture exposure or vehicle (F0 generation). (**a-d left**) Cumulative percent day at vaginal opening (VO) of rats exposed to the mixture of EDC *in utero* and through lactation (F1 generation, n=51-56/group), through germ-cell (F2 generation, n=50-52/group) or not being directly exposed (F3 generation, n=15-24/group; F4 generation, n=47-64/group). (**a and c, right**) GnRH interpulse interval measured in F1 and F3 generation females at P20 *ex vivo* through an hypothalamic explant incubation carrying out sequential sampling every 7.5 minutes for 4 hours in MEM followed by radioimmunoassay (n=4/group). Bars represent mean ± s.e.m. (*P < 0.05, ***P < 0.001 vs. CTL, Student’s t-test. Statistical comparison of vaginal opening were carried out in mean average).

**Figure 3.**
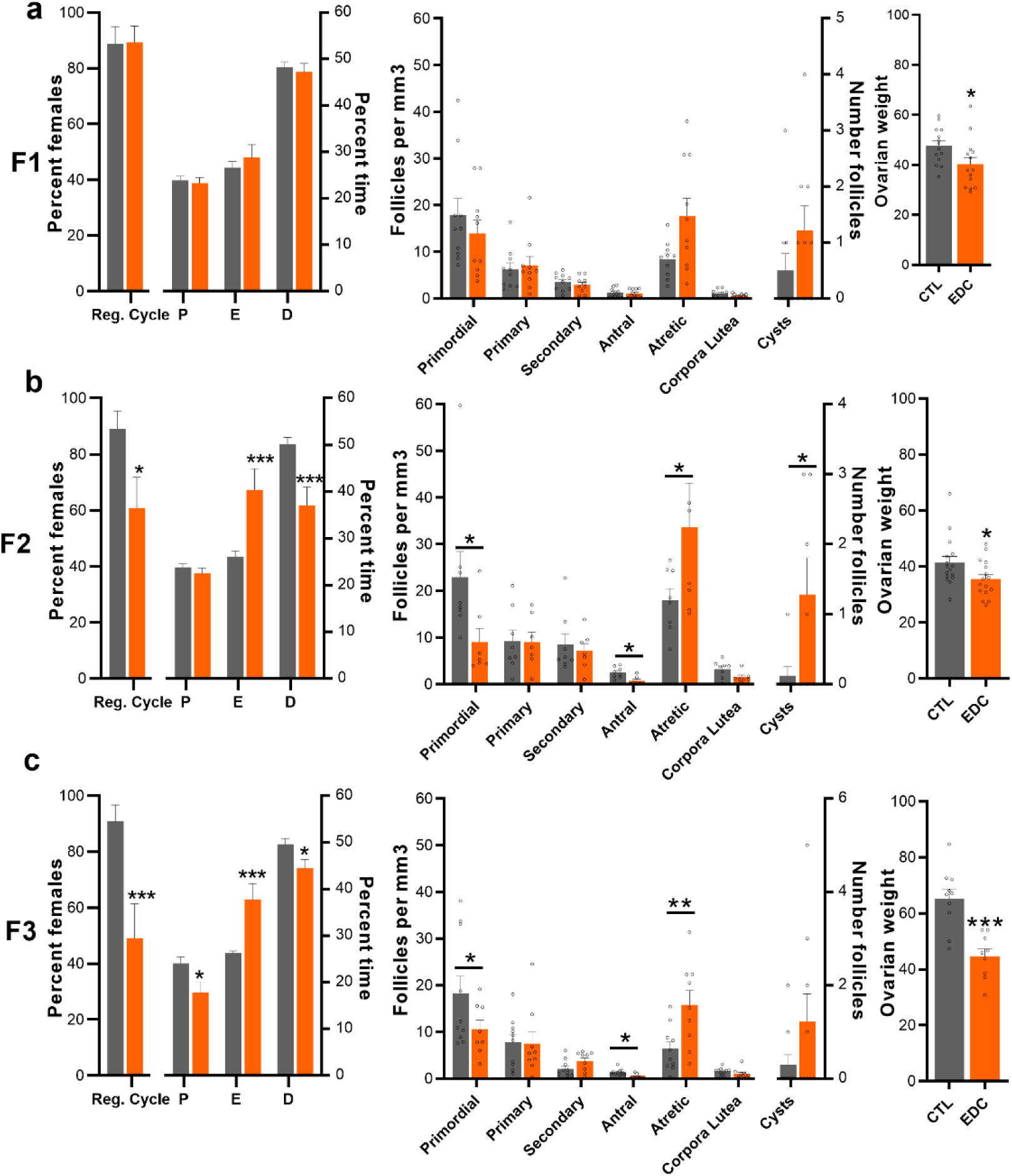
Estrous cycle, ovarian follicle development and ovarian weight across generation (F1-F3 generation) of female rats exposed to an EDC mixture or vehicle (F0 generation). (**a-c left**) Percentage of females having regular cycle and average time spent in different stages of the estrous cycle by rats exposed to the EDC mixture *in utero* (F1, n=20/group), through germ cell (F2, n=15/group) or ancestrally (F3, n=14-15/group). (**a-c middle**) Number of ovarian follicles per mm^3^ throughout development, corporal lutea and cysts quantified at P60 in F1, F2 and F3 females (n=9-10/group). (**a-c right**) Ovarian weight measured at P60 in F1 (n=14/group), F2 (n=16/group) and F3 (n=9-10/group) females. Reg. cycle= regular cycle, P= proestrus, E= estrus, D= diestrus. Bars represent mean ±s.e.m. (*P < 0.05, **P < 0.01, ***P < 0.001 vs. CTL, Student’s t-test or two-way ANOVA).

At P70, ovaries from F1, F2 and F3 females were evaluated for ovarian folliculogenesis. *In utero* exposure to EDC did not affect follicular development in F1-EDC females (Fig. 3a right). In contrast, F2-EDC and F3-EDC females displayed a significant decrease in antral follicles and an increase in atretic follicles (Fig. 3b-c right) as compared to controls. Additionally, F2-EDC females showed a significant decrease in the number of primordial follicles (Fig. 3b right). Ancestral exposure to the EDC mixture also decreased ovarian weight over time and increase the number of cysts.

Altogether, these results indicate that the EDC mixture exposure did not affect pubertal onset or ovulation in F1 females, a generation directly exposed *in utero* during development, while the next generations (F2, exposed as primordial cells in the ovaries of the F1) and F3 and F4 (ancestrally exposed to EDC) displayed delayed maturation of GnRH secretion, delayed pubertal onset and abnormal folliculogenesis.

### EDC mixture exposure disrupts the epigenetic programming of the hypothalamus

We hypothesized that the delay in GnRH secretion maturation and pubertal onset observed in F3-EDC females could be due to transcriptional and epigenetic disruption of the hypothalamic networks controlling pubertal development. Next, we used massive parallel RNA sequencing to identify gene regulatory pathways altered by EDC exposure in the MBH of F3-EDC females at P21, corresponding to the juvenile phase of prepubertal development. Gene ontology analysis showed that the main downregulated biological processes were: Neurohypophyseal hormone activity, Neuropeptide receptor activity and Dopamine signaling. Some of the downregulated molecular functions affected included: GABA, Dopamine and Glutamate signaling, while Xenobiotic processes were upregulated (Supplementary Fig. 1). qPCR quantification of hypothalamic genes critical for pubertal onset validated the downregulation of *Kiss1, Esr1*, and *Oxt* in F3-EDC females at P21 and P60 (Fig. 4a). We also identified a set of genes involved in the control of energy balance and reproduction (*Cart, Pomc, Grin2d, Gri2 and Avp*) and stress responsiveness (*Nr3c1* and *Crh*) that were altered at P21 in F3-EDC females as compared to controls (Fig. 4a, Supplementary Fig. 2).

**Figure 4.**
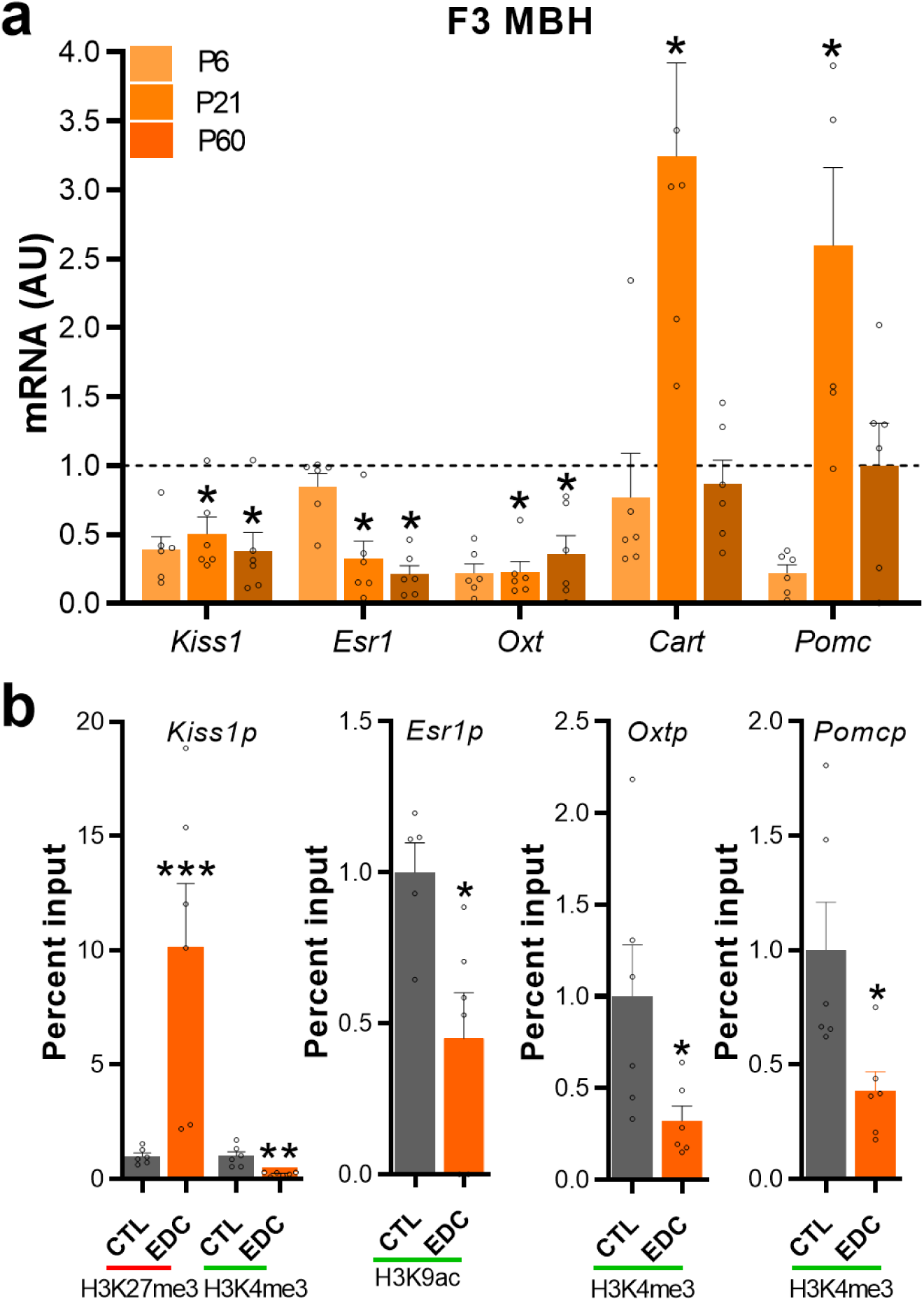
*Kiss1, Esr1, Oxt, Cart* and *Pomc* mRNA expression and promoter chromatin state in the female rat ancestrally (F3 generation) exposed to an EDC mixture or vehicle. (**a**) Expression of *Kiss1 Esr1, Oxt, Cart* and *Pomc* mRNA in the MBH of infant (P6), prepubertal (P21) and adult (P60) female rats as determined by qPCR (n=6/group). AU = arbitrary units. RNA expression data were normalized by dividing each individual value by the average of the control group at every time point. (**b**) Abundance of the TrxG-dependent activating marks H3K4me3 and H3K9ac and the PcG-dependent repressive mark H3K27me3 at the *Kiss1, Esr1, Oxt* and *Pomc* promoter in the prepubertal MBH of females ancestrally exposed to a mixture of EDC (F3 generation), as measured by ChIP (n=6/group). Bars represent mean ± s.e.m. (*P < 0.05, **P < 0.01, ***P < 0.001 vs. CTL, Student’s t-test).

In order to gain insight into the potential contribution of histone modifications to the transcriptional changes caused by EDC exposure, we used ChIP-assay to quantitate the repressive (H3K27me3 and H3K9me3) and activating (H3K4me3, K3K9ac) histone modifications at the promoter region of target genes in the hypothalamus of F3 control and EDC females (Fig. 4b, Supplementary Fig. 3). Transcriptional downregulation caused by EDC mixture was consistently associated with either an increase in repressive histone marks or a decrease in activating histone modifications at the promoter of genes critical for the onset of puberty. We observed an increase in the repressive marks H3K27me3 at the promoter region of the *Kiss1* gene, while a decreased abundance of activating marks H3K4me3 or K3K9ac was observed at *Esr1, Kiss1* and *Oxt* (Fig. 4b, Supplementary Fig. 3). No change in histone landscape was detected at the *Cart* promoter while a decrease in the repressive H3K9me3 was detected at the *Pomc* promoter, suggesting that the potential role of EDC exposure in the epigenetic programing of the hypothalamus is locus specific (Supplementary Fig. 3).

### EDC mixture disrupts the dopaminergic control of maternal behavior throughout generations

Phenotypic inheritance of EDC effects has been shown to be transmitted through epigenetic changes in germ-cells^16^. Alternatively, maternal behavior is known to affect pubertal development^17^ and can be passed down throughout generations by inducing epigenetic changes at the somatic level. Because EDC have been shown to affect maternal care^18^, we aimed at exploring the effects of the EDC mixture on maternal care throughout generations. Direct exposure to the EDC mixture (F0 generation) did not alter maternal care (Fig. 5a). F1, F2 and F3-EDC dams displayed impaired maternal behavior characterized by lower levels of licking and grooming compared to the control dams (Fig. 5b-d). Additionally, females exposed *in utero* (F1 generation) or as germs cells (F2 generation) spent more time resting alone or being active outside the nest, respectively. No changes were observed in retrieval, nursing, nest building or in average in-nest activity throughout generations (Supplementary Fig. 4-9).

**Figure 5.**
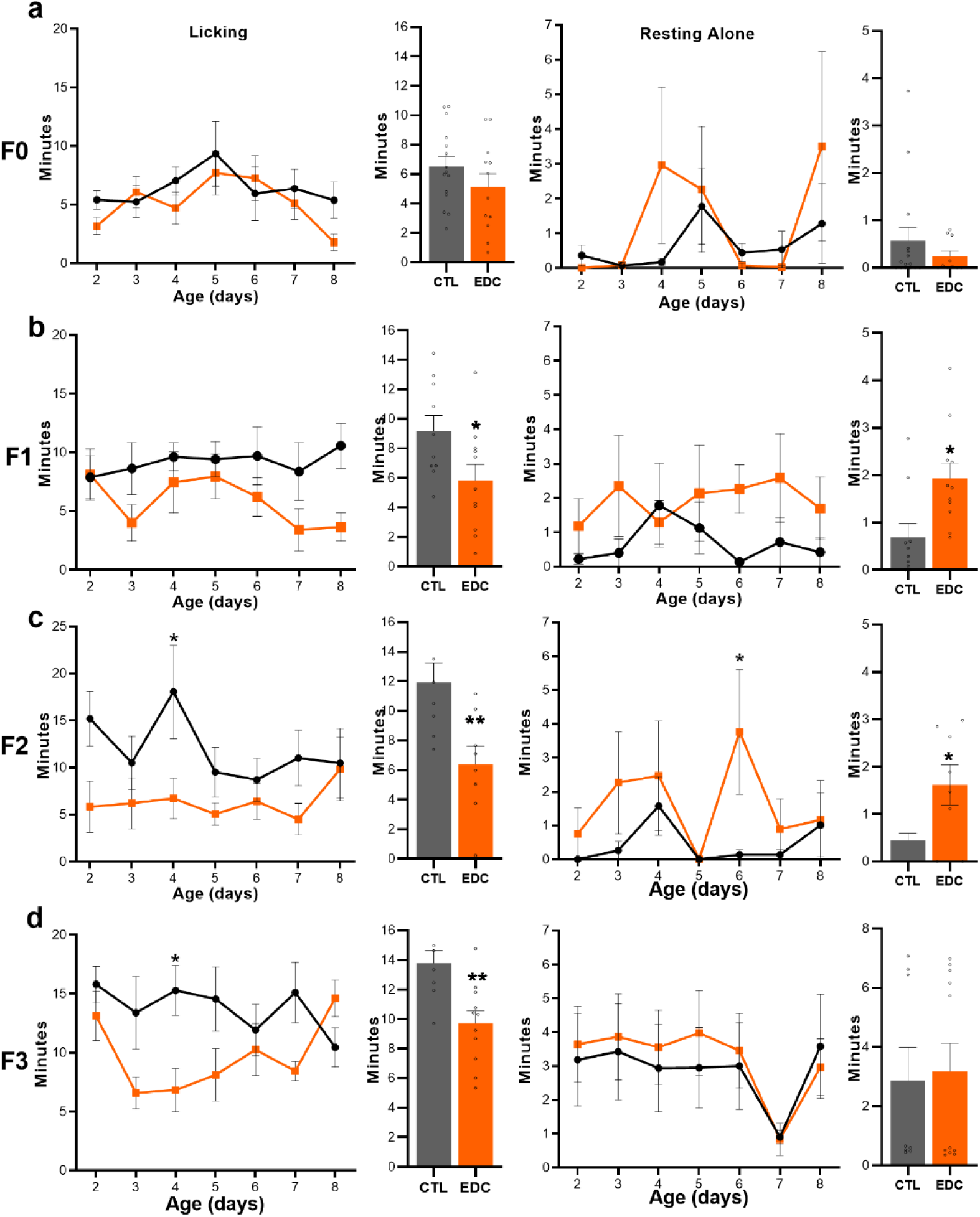
Maternal behavior displayed by female rats exposed to a mixture of EDC throughout 4 generations. (**a-d left**) Time spent by dams displaying licking and grooming behavior toward pups after direct (F0 generation, n=15/group), *in utero* and through lactation (F1 generation, n=10-11/group), germ-cell (F2 generation, n=11/group) or ancestral (F3 generation, n=11/group) exposure to an EDC mixture or vehicle from P2 to 8. Bar graph shows pooled time licking and grooming from P2-8. (**a-d right**) Time spent by dams resting alone outside the nest not being involved in maternal care throughout generations (F0-F3 generation). Bar graph shows pooled time spent resting alone from P2-8. Plotted lines represent average of time ± s.e.m. (*P < 0.05 vs. CTL, two-way ANOVA followed by Sidak’s multiple comparisons test).

In order to identify the hypothalamic targets potentially involved in the alterations of maternal care programming caused by EDC, we searched the massive parallel RNA sequencing gene ontology analysis described before in MBH of F3-EDC females (Supplementary Fig. 2) focusing on those downregulated genes that belong to the enriched categories: social behavior, maternal behavior, grooming behavior and synaptic dopaminergic transmission (Supplementary Fig. 1). Additional gene ontology analysis of massive parallel RNA sequencing done in MBH of F1 females showed downregulated categories related to D2 dopamine receptor binding, glutamate binding and neurotransmitter uptake (Supplementary Fig. 8). qPCR validation of common target genes in F1 and F3 gene ontology analysis showed that critical dopaminergic signaling (*Th, Dnm1* and *Darpp32)*, involved in maternal motivation, was significantly decreased in the hypothalamus of F1-EDC animals at P21 and/or 60 (Fig. 6a). On the other hand, the dopaminergic receptor 1 (*Drd1)* was found to be upregulated, a possible sign of a compensatory mechanism induced by reduced synaptic dopamine (Fig. 6a). Moreover, genes associated with stress responsiveness (*Nr3c1* and *Crh*) were not altered (Supplementary Fig 9a). Overall, these data indicate that exposure to EDC decreased the expression of key genes involved in the dopaminergic control of maternal behavior.

**Figure 6.**
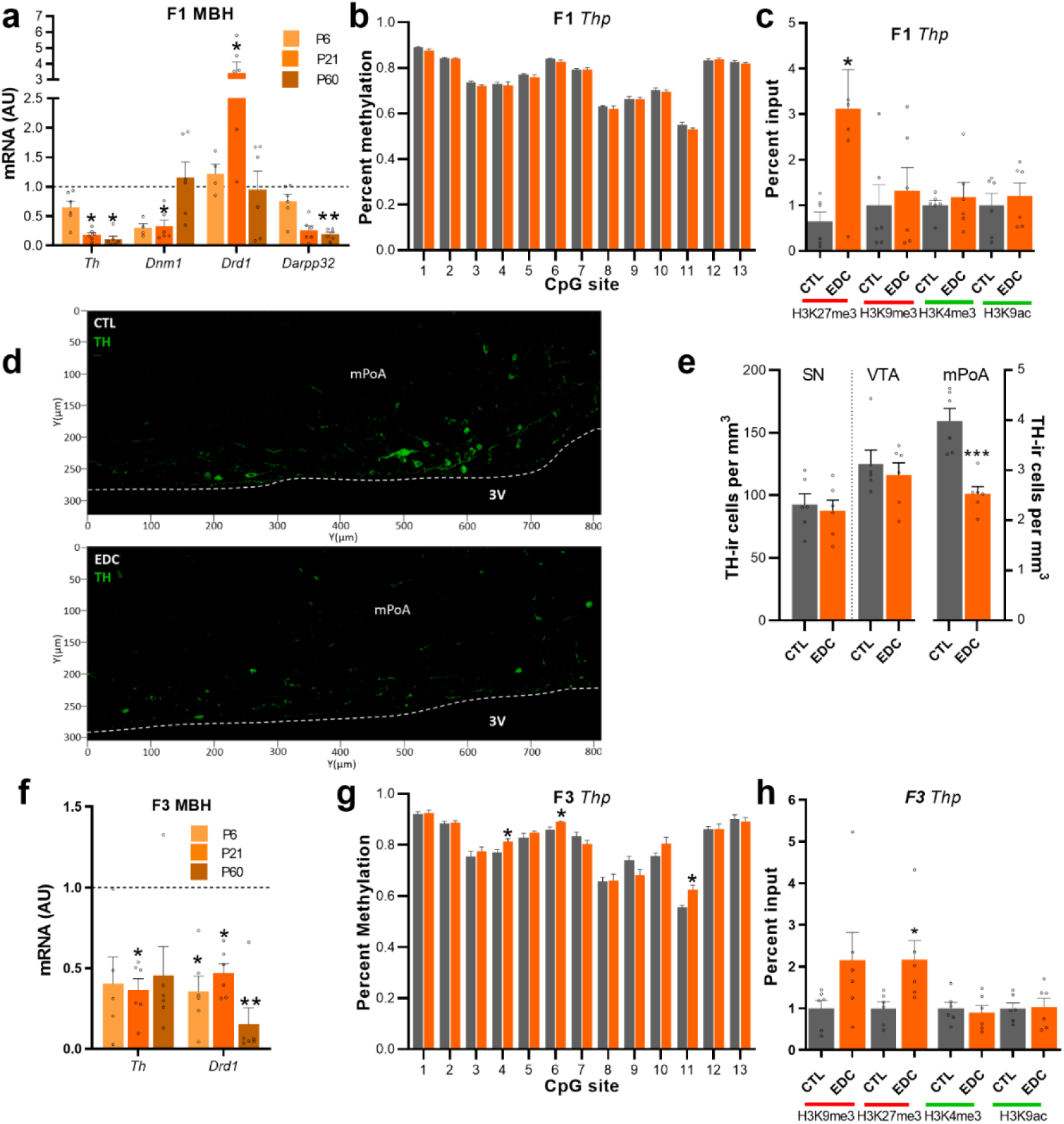
Dopaminergic signaling proteins, mRNA expression and chromatin state in the female rat *in utero* (F1 generation) and ancestrally (F3 generation) exposed to an EDC mixture or vehicle. (**a**) Expression of *Th, Dnm1, Drd1* and *Darpp32* mRNA in the MBH of infant (P6), prepubertal (P21) and adult (P60) female rats in utero and lactationally (F1 generation) exposed to an EDC mixture as determined by qPCR (n=6/group). RNA expression data were normalized by dividing each individual value by the average of the control group at every time point. (**b**) Methylation state at 13 CpG sites of the Th gene promoter from MBH explants of F1 females at P2 (n=6/group). (**c**) Abundance of the TrxG-dependent activating marks H3K4me3 and H3K9ac and the PcG-dependent repressive mark H3K27me3 and H3K9me3 at the *Th* promoter in the prepubertal MBH of females EDC and control from the F1 generation, as measured by ChIP (n=6/group). (**d**) Abundance of Th-ir cells (green color) within the mPoA of prepubertal female rats. 3V, third ventricle. (**e**) Quantification of Th immunoreactivity in the SN, VTA and mPoA of females from the F1 generation. Bars represent mean of cells per mm^3^ ± s.e.m; (**f**) Expression of *Th* and *Drd1* mRNA in the MBH of infant (P6), prepubertal (P21) and adult (P60) female rats ancestrally (F3 generation) exposed to EDC mixture (n=6/group). (**g**) Methylation state of at 13 CpG sites of the Th gene promoter from MBH explants of F3 females at P2 (n=6/group) (**h**) Abundance of H3K4me3,H3K9ac, H3K27me3 and H3K9me3 histone posttranslational modifications at the *Th* promoter in the prepubertal MBH of females EDC and control from the F3 generation, as measured by ChIP (n=6/group). AU = arbitrary units. Bars represent mean ± s.e.m. (*P < 0.05, **P < 0.01, ***P < 0.001 vs. CTL, Student’s t-test).

Epigenetic analysis of the *Th* promoter (*Thp*) region identified no change in DNA methylation (Fig. 6b), but higher levels of the repressive histone H3K27me3 modification in the hypothalamus of F1-EDC females (Fig. 6c) without affecting H3K9me3, H3K4me3 or H3K9ac status. No alterations in histone modification landscape were detected in the 5’ regulatory regions of *Darpp32* or *Drd1* (Supplementary Fig. 9b), suggesting that transcriptional alterations caused by EDC do not involve epigenetic reprogramming in the promoter region of those genes and demonstrating that the epigenetic alterations are targeted to specific loci.

To further characterize the loss in dopaminergic signaling, we quantified the TH-immunoreactivity in the substantia nigra (SN), the ventral tegmental area (VTA) and the median preoptic area (mPoA), part of the VTA-mPOA-NAc pathway involved in maternal motivation^19^. Exposure to EDC significantly decreased the number of Th immunopositive cells in the mPOA of F1 females at P21 (Fig. 6d-e), confirming the mRNA expression results. No differences were observed in the VTA or NAc.

Because the F3 generation also showed diminished licking and grooming behavior as a result of the EDC mixture treatment of the F0 generation, we hypothesized that the dopaminergic system would also be affected. *Th* and *Drd1* mRNA expression were found to be significantly reduced in hypothalamus of F3-EDC females (Fig. 6f). *Th* downregulation was associated with increased DNA methylation at 3 specific CpGs in the *Th* promoter, as well as enhanced repressive H3K27me3 and H3K9me3 histone marks (Fig. 6g-h). No differences were found in the activating histone modifications H3K4me3 and H3K9ac. These results indicate that the transgenerational reduction of licking and grooming behavior is also linked to diminished hypothalamic dopaminergic signaling by increased action of the Polycomb group (PcG) of epigenetic repressors, and maybe other histone methyl transferases on the *Th* promoter.

So far, our data show a direct effect of the EDC mixture on the prenatal development of the F1 generation’s dopaminergic system throughout diminished dopamine signaling and reduced maternal behavior. The perpetuation of this phenotypic manifestation throughout generations, even in the absence of EDC exposure, is also associated to hypothalamic dopaminergic loss, but through epigenetic reprograming mediated by the PcG and increased DNA methylation at specific CpGs in the *Th* promotor.

### Sexual maturation is altered through germ-cell transmission

Our data identified a multigenerational transmission of maternal care starting with the F1 all the way down to the F3 generation. Sexual maturation was found to be delayed in F2 to F4 generations and associated with epigenetic reprograming of the hypothalamus identified in the F3 generation. As no differences in sexual maturation was found in *in utero* exposed females (F1 generation), we hypothesized that the delay in sexual maturation in F2 and consecutive generations could be explained by hypothalamic reprograming caused by variations in maternal care. To determine whether delayed puberty is caused by germ-cell or experience-based inheritance, a cross-fostering paradigm was carried out. Control F2 pups raised by CTL (CC) or EDC (EC) F1 dams showed normal vaginal opening and estrous cyclicity. To the contrary, germ-cell EDC exposed pups (F2 generation) raised by an EDC (EE) or CTL (CE) dams, indistinctively showed a delay in vaginal opening (Fig. 7a) and disrupted estrous cycles, characterized by increased time spent in estrus and decreased time spent in diestrus (Fig. 7b). F2 germ-cell exposed pups showed a downregulation in the hypothalamic expression of *Kiss1, Esr1* and *Oxt* at P21, independently of being raised by an EDC (EE) or control (CE) dam and these changes persisted through the next generation (Fig. 7c-f). Downregulation of *Kiss1, Esr1* and *Oxt* was consistently associated with abundance of histone marks, showing a repressive state (Fig. 7g-I, Supplementary Fig. 10). These results show that cross-fostering could not restore normal pubertal timing in germ-cell EDC exposed animals, nor did EDC exposed dams cause delayed sexual maturation of unexposed pups. Our results demonstrate that delayed sexual maturation is caused by a germ-cell inheritance of EDC mixture effects.

**Figure 7.**
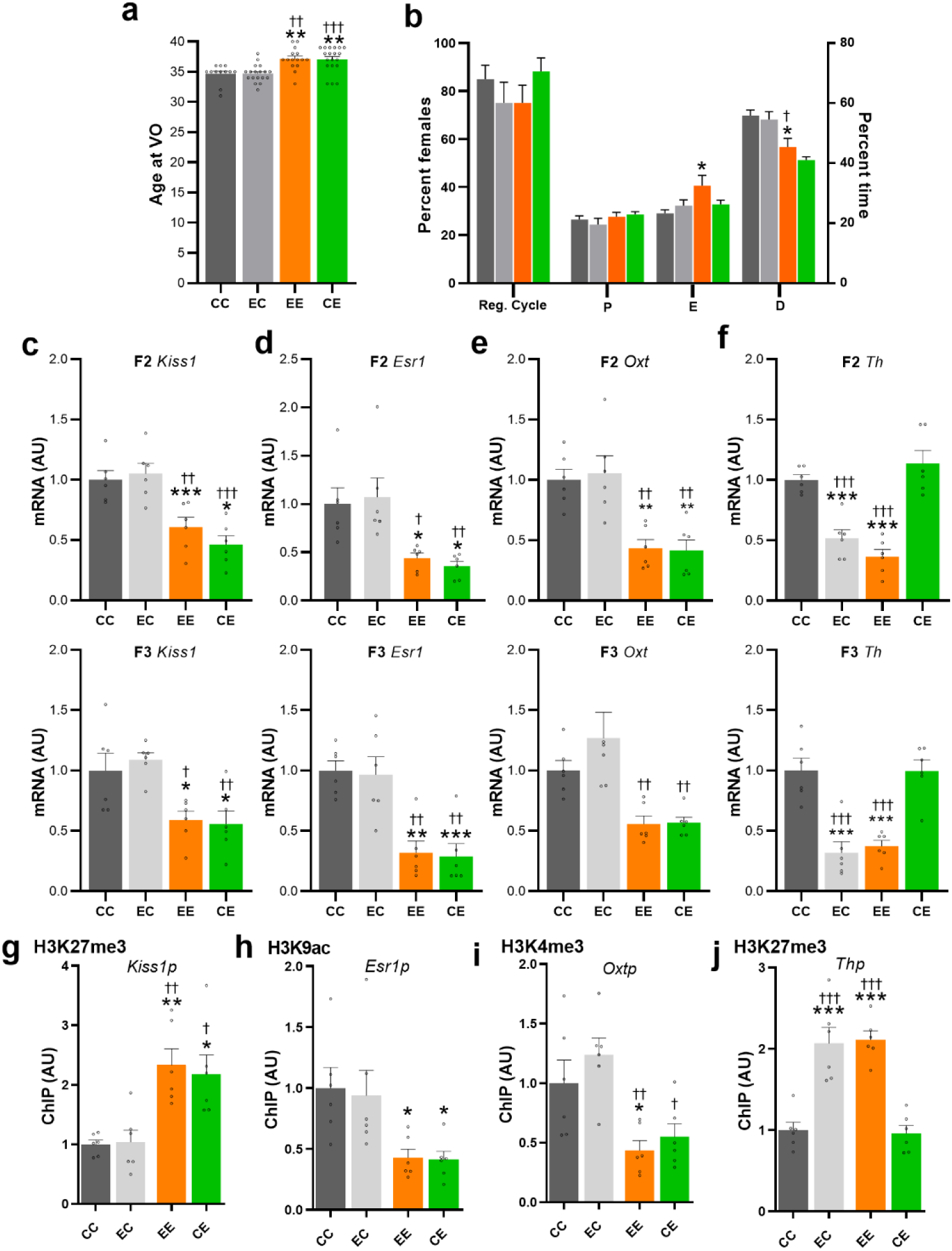
F2 cross-fostered offspring sexual maturation and mRNA expression data. (**a**) Average age at vaginal opening of cross-fostered germ-cell EDC exposed pups or control raised by either *in utero* EDC exposed dams or control (n=12-19/group) (**b**) Percentage of cross-fostered females having regular cycle and time spent in different stages of the estrous cycle (n=10/group) (**c-f**) Expression of *Kiss1 Esr1, Oxt*, and *Th* mRNA in the MBH of F2 and F3 generation crossfostered pups at P21, as determined by qPCR (n=6/group). (**g-j**) Abundance of the TrxG-dependent activating marks H3K4me3, H3K9ac or the PcG-dependent repressive mark H3K27me3 at the *Kiss1, Esr1, Oxt* and *Th* promoter in the prepubertal MBH of females crossfostered from the F3 generation, as measured by ChIP (n=6/group). Reg. cycle= regular cycle, P= proestrus, E= estrus, D= diestrus. CC= control pup raised by control dam; EC: control pup raised by *in utero* EDC exposed dam; EE= germ-cell EDC exposed pup raised raised by *in utero* EDC exposed dam; CE= germ-cell EDC exposed pup raised by control dam. Bars represent mean ± s.e.m. (*P < 0.05, **P < 0.01, ***P < 0.001 vs. CC; †P < 0.05, ††P < 0.01, †††P < 0.001 vs. CE, one-way ANOVA).

Here we also show that in F1 females, the dopaminergic system abnormalities are associated with alterations in maternal care. Our cross-fostering paradigm reveal that *Th* mRNA levels are downregulated in F2 and F3 generations when raised by an EDC-exposed mother, independently of their ancestral as germ cells (EE or EC) exposure, suggesting that the alterations in the dopaminergic system are transmitted through maternal care (Fig. 7f). *Th* downregulation was associated with enhanced repressive H3K27me3 histone marks as found in the previous experiment (Fig. 7j, Supplementary Fig. 10).

Overall, our results demonstrate that direct EDC mixture action on F1 females during development diminishes hypothalamic dopaminergic signaling impacting licking and grooming behavior. This behavior is “learned” as it seems to be transmitted down to the next generations. On the other hand, the delayed sexual maturation is transmitted throughout generations and cannot be corrected when pups are cross-fostered with controls dams. These results suggest a germline transmission of the reproductive phenotype.

## DISCUSSION

The potential contribution of developmental exposure to a low dose EDC mixture in the neuroendocrine regulation of female reproductive and maternal behaviors, to the best of our knowledge, has never been addressed. In the present report, we provide evidence that EDC exposure induces multi- and trans-generational alterations of maternal care and sexual maturation, respectively, throughout epigenetic reprogramming of the hypothalamus. Our results identify a delay in pubertal development that appears apparent starting at the F2 generation, in females exposed to the EDC mixture through the germline. The reproductive phenotype is found up to the F4 generation and is determined as a delay in the day of vaginal opening and increase in the GnRH interpulse interval. These animals showed disrupted estrous cyclicity, with increased time in estrus and decrease time in proestrus and diestrus, a sign of diminished ovulatory cycle efficiency. Moreover, EDC exposed animals showed a significant reduction in antral and enhanced atretic follicles, a clear sign of subfertility. Surprisingly, no reproductive effects were detected in the F1generation, the one directly exposed to the EDC mixture during embryonic and lactational development, which suggests that the reproductive phenotype appears as a consequence of germline exposure.

The activation of the GnRH network during the juvenile period involves the transcriptional activation of KNDy neurons, those that co-express Kisspeptin, Neurokinin-b and Dynorphin. We have recently deciphered that female sexual maturation is controlled by the dual coordinated action of the Polycomb (PcG) and Trithorax (TrxG) group of chromatin remodelers, leading to a concomitant epigenetic and transcriptional switch from repression to activation of the *Kiss1* locus in KNDy neurons of the ARC^13,14^. Moreover, we recently described that metabolic cues target KNDy neurons through epigenetic modifications affecting pubertal timing^20^. Specifically, undernutrition-induced delayed pubertal onset was found to be related to a developmental increase in the repressive histone deacetylase Sirtuin 1(SIRT1) responsible for decreased H3K9/16ac at the Kiss1 gene promoter, as well as an increase in H3K27me3, a repressive histone modification related to the activity of the PcG of epigenetic repressors. Since developmental exposure to the plasticizer BPA can disrupt kisspeptin neurons in adults^21,22^, we hypothesized that one putative target of EDC mixture induced disruption of puberty could be the KNDy neuron. RNAseq analysis of the hypothalamus of F3 females ancestrally exposed to the EDC mixture demonstrated that the reproductive effects where associated with decreased *Kiss1, Oxt* and *Esr1* expression, three main players in the control of the GnRH pulse generator and the correct timing of puberty and ovulation^23–27^. Here we found that F3 descendants of EDC treated animals show a disbalanced histone configuration at the *Kiss1* promoter with increased repressive H3K27me3 and reduced H3K4me3, a histone modification found at promoter regions of activated genes, suggesting a central role of the PcG/TrxG balance in the EDC induced delay of puberty. Moreover, although we found that the hypothalamic expression of *Pomc* and *Cart*, two postulated metabolic activators of the GnRH system^28,29^, is transiently increased, it did not overcome the effects of the *Kiss1, Esr1* and *Oxt* loss on pubertal delay. The loss in *Esr1* expression was associated with the diminished presence of H3K9ac at its promoter region, suggesting a role of the sirtuin deacetylases as putative regulators of EDC mediated reproductive effects. Our results suggest that EDC mixture exposure affects the epigenetic programming on the hypothalamus throughout generations delaying sexual maturation and altering estrous cyclicity and folliculogenesis. A similar transgenerational phenotype was observed after developmental exposure to pesticides, jet fuel, DEHP and TCDD, displaying altered puberty, lower primordial follicles and accelerated follicle recruitment^6,30–32^. Furthermore, there are striking similarities between the epigenetic changes involved in the EDC induced reprograming of the hypothalamus and those induced by undernutrition. In both cases we identified a predominant role of the PcG and sirtuins in repressing gene expression by increasing the repressive role of H3K27me3 and decreasing the effects of H3K9ac at the promoter region of puberty activating. This observation is in line with several reports suggesting that EDC function as metabolic disruptors by sharing common intracellular pathways^33^.

It is known that alterations in maternal care are transmitted through generations affecting the hypothalamic response to stress^34^ and also impacting reproductive development by modulating *Esr1* expression^35^. Our ancestrally EDC mixture exposed (F2-F3) animals showed delayed puberty of central origin and diminished pup licking behavior, without affecting any other maternal behavior. Surprisingly, the F1 generation, that had normal pubertal development and ovarian phenotype, also showed diminished pup licking behavior, suggesting that direct EDC mixture exposure during intrauterine development directly affects the neuronal network involved in parental behavior. Gene ontology analysis of hypothalamic RNAseq demonstrated that the dopaminergic system is one of the main targets of the EDC mixture in the F1 generation. In particular, EDC exposed F1 females showed a significant loss in *Th* expression accompanied with increased repressive H3K27me3, a main target of PcG action. Moreover, this effect is localized to the mPoA, since there was no TH staining loss in either the SN or the VTA, demonstrating that the action of the EDC mixture is region specific and directed to epigenetically reprogram the dopaminergic system. On the other hand, females of the F3 generation also showed diminished dopaminergic signaling in the hypothalamus, but with further penetrance of the epigenetic phenotype. In this case, the *Th* promoter not only showed increased repressive H3K27me3 but also H3k9me3, as well as increased CpG methylation at 3 independent sites throughout the *Th* promoter. This shows that the epigenetic reprograming of the *Th* locus of the F3 generation differs of that of the F1, since the F3 generation was not directly exposed to the EDC mixture but it was exposed to a dysfunction in maternal care, raising the possibility of direct multigenerational transmission of the behavioral phenotype. Such alteration is multigenerationally transmitted through epigenetic alteration of the dopamine system.

In the current study, we have found a transcriptional and epigenetic alteration of stress response genes in F3 females raised under diminished maternal care. F1-EDC females did not show differences in hypothalamic *Nr3c1* or *Crh* expression, while receiving normal maternal care from their F0 mothers. This confirms previous studies showing that maternal care induces increased responses of the HPA axis, decreasing *Nr3c1* transcriptional levels via epigenetic alterations throughout generations^36,37^.

We also identified that the reproductive phenotype is transmitted through the germ line and is not explained by impaired maternal behavior, since the crossbreeding of the F2 generation with non-exposed mothers did not normalize pubertal timing nor the downregulation of hypothalamic *Kiss1, Esr1* and *Oxt* expression in the F2 or F3 generations. These effects are not general since the cross-fostered EDC exposed animals raised by control females showed a normalization of the hypothalamic dopaminergic network by increased *Th* expression, indicating that the reduction in maternal behavior is in fact learned and multigenerationally transmitted from the F1 generation. The transgenerational disruption of sexual maturation phenotype could be caused by germ cell alterations of epigenetic developmental programs induced by EDC treatment. This EDC-induced epigenetic reprogramming needs to be resistant to erasure and to be transmitted across multiple generations^38^. Thereafter, EDC-induced germ cell alterations flows from germline to soma^39^, affecting in this case the organization of the GnRH network. It is possible that environmental factors alter the germline epigenome directly or indirectly through soma-to-germline transmission. For instance, it has been recently reported that small RNA molecules are transferred from somatic cells to germ cells, and that EDC treatment affects ncRNAs in germ cells ^40–42^. Further studies should address these issues and determine the developmental window at which the germ cells epigenome is more sensitive to environmental alterations.

In current human pregnancy conditions, virtually any woman and her fetus are exposed to a low-dose mixture of at least 100 EDC^1^. Small-scale studies with women visiting fertility clinics show that human follicular fluid samples contain a wide array of chemicals with endocrine disrupting activity, such as DDT, phthalates, bisphenol A and perfluorinated compounds, indicating direct exposure of maturing oocytes and their surrounding steroid-producing cells^43–45^. Most studies are not designed to investigate environmentally relevant combinations. However, such studies are crucial from a regulatory point of view because current risk assessment is based on the effects of individual chemicals. In order to address the issue of complex mixture, we have selected chemicals for which rudimentary information about their endocrine disrupting effects *in vivo* and data about human exposures were available to guide the choice of doses^46^. Developmental exposure to this mixture at high doses has been previously shown to alter sexual differentiation in males^47,48^ and disrupts estrous cyclicity in females when exposed *in utero*^49,50^. However, this study is the first one to our knowledge to report trans- and multi-generational effects of a low dose mixture on the hypothalamic control of puberty and maternal behavior.

Altogether, the present results demonstrate that developmental exposure to a human relevant dose of an EDC mixture alters the hypothalamic epigenetic programming of reproduction trough increased action of the PcG and possibly sirtuins on key puberty activating genes and that these effects are trans-generationally transmitted. In addition, direct exposure to the EDC mixture downregulates the hypothalamic dopaminergic network by increased action of the PcG and other methyltransferases, affecting maternal behavior in a multigenerational and reversible manner.

## MATERIAL AND METHODS

### Animals

Adult Female Wistar rats purchased from the animal facility of the University of Liège were housed individually in standardized conditions (12h inverted dark/light phase, 22.8°C and food and water *ad libitum*). All animals were raised in EDC-free cages (Polypropylene cages, Ref 1291H006, Tecnilab, Netherlands) and fed EDC- and phytoestrogen-free chow (V135 R/Z low phytoestrogen pellets, SSNIFF Diet, Netherlands). Water was supplied in glass bottles. All experiments were carried out with the approval of the Belgian Ministry of Agriculture and the Ethics Committee at the University of Liege.

### Chemicals

The endocrine disrupting chemicals di-n-butylphthalate (DBP) (purity >99.0 %, 84-74-2), di-(2-ethylhexyl)phthalate (DEHP) (purity >99.5 %, 117-81-7), vinclozolin (purity >99.5 %, 50471-44-8), prochloraz (purity >98.5 %, 67747-09-5), procymidone (purity >99.5 %, 32809-16-8), linuron (purity >99.0 %, 330-55-2), epoxiconazole (purity >99.0 %, 106325-08-8), 2-ethylhexyl 4-methoxycinnamate (OMC, EHMC) (purity >98.0 %, 5466-77-3), dichlorodiphenyldichloroethylene (p,p’-DDE) (purity >98.5 %, 72-55-9), 4-methyl-benzylidene camphor (4-MBC) (purity >98.0 %, 36861-47-9) and butylparaben (purity >99.0 %, 94-26-8) were purchased from AccuStandard. Bisphenol A (purity >99.5 %, 80-05-7), and paracetamol (purity >99.0 %, 103-90-2) and corn oil (as a control vehicle) were obtained from Sigma-Aldrich. EDC compounds were dissolved in corn oil in order to obtain the final concentration showed in Table 1.

**Table 1.** Features of the 13 chemicals included in the EDC mixture

| Substance | Function | AA/E | Dose<br>($\mu\text{g}/\text{kg}/\text{d}$) | NOAEL<br>($\text{mg}/\text{kg}/\text{d}$) | LOAEL |
| --- | --- | --- | --- | --- | --- |
| Di-n-butyl-phthalate (DBP) | Plasticizers | AA | 10 | 50 | 100 |
| Di-(2-ethylexyl)-phthalate (DEHP) |  | AA | 20 | 3 | 10 |
| Bisphenol A (BPA) |  | AA / E | 1,5 | 5 | 1,2 |
| Vinclozolin | Dicarboximide fungicide | AA | 9 | 5 | 10 |
| Procymidon |  | AA | 15 | 10 | 25 |
| Prochloraz | Imidazole fungicide | AA | 14 | 5 | 10 |
| Linuron | Urea-based herbicide | AA | 0,6 | 25 | 50 |
| Epoxinaxole | Triazole fungicide | AA | 10 | 15 | - |
| 4-Methyl-benzylidene camphor (4-MBC) | UV-filter | E | 60 | 0,7 | 7 |
| Octyl methoxycinnamate |  | E | 120 | - | 500 |
| p'p'-DDE | Metabolite of insecticide DDT | AA / E | 1 | - | 10 |
| Butyl paraben (BP) | Antifungal and antibacterial preservative | AA | 60 | - | 100 |
| Acetaminophen | Analgesic & antipyretic | AA | 800 | - | 350 |

**Notes:** AA: antiandrogenic, E: estrogenic, NOAEL: non-observed adverse effect level, LOAEL: low-observed adverse effect level

### Experimental design

After habituation to the animal care facility, F0 generation females were mated for two weeks with a wild-type male in order to generate the F1 generation. F0 females were exposed to an EDC mixture or corn oil (vehicle) from two weeks before gestation to the last day of lactation (Fig. 1). Daily exposure was done by injecting 50µl of EDC mixture or corn oil in a waffle and allowing them to eat it. Verification was systematically done after 10 minutes of exposure. Females were randomly assigned to treatment. At postnatal day (P) 70, *in utero* EDC mixture exposed and control females from the F1 generation were mated with a wild-type male to generate the F2 generation. Similar procedure was done to generate the F3 and F4 generation. Pups from every generation were homogenized by sex-ratio and litter size and followed for developmental weight until weaning (P21). From weaning to P70, control females were paired with a female EDC and housed together under the same conditions. Behavioral and reproductive data were obtained from two independent non-related cohorts of animals.

### Cross-fostering

To distinguish between germ-cell versus experience-based phenotype transmission, a cross-fostering paradigm was used. In order to avoid an effect of cross-fostering, a maximum of two pups per litter were cross-fostered in pups from F1 female dams as followed. F1 generation control dams raised either EDC mixture exposed pups (CE) or control pups from another dam (CC). F1 generation EDC mixture exposed dams raised either EDC mixture exposed pups from another dam (EE) or control pups (EC). Cross-fostering was carried out within the first 24h after delivery and a 1mm tail incision was done in order to distinguish cross-fostered pups.

### Maternal Behavior

To assess the effect of an EDC mixture exposure throughout generations on maternal behavior, a set of maternal behaviors were quantified in EDC mixture and control lactating females from the F0 to the F3 generation. Additionally, we have quantified maternal behavior in F1 dams (EDC mixture exposed or control dam with either control or EDC mixture exposed pups from another dam) with cross-fostered pups and in F2 cross-fostered lactating females (EDC mixture exposed or non-exposed females raised by an EDC mixture exposed or control dam). Maternal behavior was recorded with an infrared camera (Bell and Howell, DNV16HDZ-BK) from lactational day 2 to 8 for 1 hour during the dark phase. Randomization was done in order to avoid recording every female at the same hour of the day. A set of in-nest behaviors (retrieval, mouthing, licking/grooming, arched-back/blanked/passive nursing and nest building) and off-nest behaviors (eating/drinking, grooming, active or resting alone) were quantified as reviewed in^51^ by an experimenter blind to condition.

### Pubertal onset and estrous cyclicity

To assess the effect of an EDC mixture exposure throughout generation on sexual maturation, females from the F1 to the F4 generation were followed for vaginal opening and estrus cyclicity as described previously^13,52^. Briefly, from P25, females were daily inspected for vaginal opening by two experimenters. From the day of vaginal opening to P70 estrous cycle was evaluated with vaginal smears that were taken every day during the morning before the lights off at 4pm. Regular cycles were defined as a sequence of diestrus 1, diestrus 2, proestrus and estrus in 4 consecutive days^53^. The percentage of females having a regular cycle and the time spent in every stage of the cycle were calculated from week 6 to week 10 of age.

### Ovarian histology and uterus weight

To assess the effect of an EDC mixture exposure on folliculogenesis throughout generations, ovaries from F1 to F3 generation females at P70 were removed for histological quantification of follicle development. After removal, ovaries were weighted together with the uterus and fixed overnight in 4% paraformaldehyde, dehydrated in 70% EtOH and paraffin-embedded. Histological analysis was done in 8-µm coronal sections (microtome RM2245, Leica), after deparafinization and stain with hematoxylin and eosin. For quantification, every other section throughout the whole ovary were digitalized using an automated digital microscopy system DotSlide (Olympus, BX51TF, Aartselaar, Belgium). Dotslide images taken at a magnification of 10x, which were in a proprietary format were converted into a standard TIFF format and 3-fold decimated, easier to handle. Thereafter, follicles at every stage of folliculogenesis (primordial, primary, secondary, antral and atretic), cystic follicles and corpora lutea were manually quantified avoiding double-counting by an experimenter blindness to treatment with Aperio ImageScope v12.3.2.8013 software (SCR_014311, Leica Biosystems). Total ovarian volume was automatically calculated using an original program developed using the image analysis toolbox of the MatLab (SCR_001622, 2016a, The Mathworks Inc., Natick, MA, USA) software. The follicles were classified according to well-established criteria^54,55^. Double counting of late-stage follicles was avoided by digitally marking each follicle throughout the consecutive images. Each follicle was counted once whenever the oocyte was present. As analysis were done in every other section, we apply a two-fold correction factor for quantification of early stage follicles (primordial and primary follicles) to compensate sections not analysed. Measurements are expressed as number of follicles or corpora lutea per volume (mm^3^).

### Hypothalamic explants incubation and GnRH assay

To assess the effect of an EDC mixture exposure on juvenile GnRH frequency, GnRH interpulse interval was measured using a hypothalamic explants incubation system followed by a GnRH assay from prepubertal females of the F1 and F3 generation, as described previously^15,56^. Briefly, after decapitation, brain was dissected by performing two sagittal incisions along the lateral hypothalamic sulci and two transversal incisions of 2mm ahead from the anterior boundaries of the optic chiasm and along the caudal margin of the mammillary bodies. Once dissected, medial basal hypothalamus (MBH) and preoptic area (PoA) explants were transferred into an individual chamber, in a static incubator, submerged in MEM. Incubation medium was then collected and renewed every 7.5 min for a period of 4 hours. The GnRH released into the incubation medium was measured in duplicate using a radioimmunoassay method with intra and inter-assay coefficients of variation of 14 and 18% respectively. The highly specific CR11-B81 (AB_2687904) rabbit anti-GnRH antiserum (final dilution 1:80,000) was kindly provided by Dr. V.D. Ramirez (Urbana, IL)^57^. Data below the limit of detection (5 pg/7.5-min fraction) were assigned that value.

### DNA and RNA extraction, reverse transcription and RT-PCR

Expression of genes involved in the hypothalamic control of puberty, reproduction and maternal behavior were studied by quantitative PCR (qPCR) analysis using the half of MBH and mPoA explants from females exposed and non-exposed to an EDC mixture from the F1 and F3 generations at different time points (P6, P21 and P60). After decapitation, the mPoA and the MBH were rapidly dissected as described in the previous section. Additionally, MBH and PoA were divided in two by sectioning along the interhemispheric fissure. MBH fragments contains the entire arcuate nucleus. Total RNA and DNA were extracted from the half-MBH and half-mPoA tissue using All Prep DNA/RNA Mini kit (Qiagen, Germantown, MD) following the manufacturer’s instructions. Five hundred ng of RNA for each sample were reverse transcribed using the Transcriptor first strand cDNA synthesis kit (Roche, Germany). For real-time quantitative PCR reactions, the cDNA of our samples were diluted 10 fold and 4 µl were added to a mix of 5 µl FastStart Universal SYBR Green Master (Roche, Germany), 0.4 µl of nuclease-free water and 0.3 µl of forward and reverse primer (see primer sequences in Supplementary Table 1). The samples were run in triplicate using a LightCycler 480 thermocycler (Roche, Germany). Ct values were obtained from each individual amplification curve and the average Ct was calculated for each target gene in each sample. Quantification of relative gene expression was performed using the ΔΔCt method implemented with the Pfaffl equation which takes into account reaction efficiency depending on primers^58^ All assays had efficiencies between 1.9 and 2.1. β-actin was used as housekeeping gene.

### Bisulfite sequencing

Genomic DNA was Bisulfite-converted (BC) using the EZ DNA Methylation-Gold kit (Zymo Research, Irvine, CA) according to the manufacturer’s instructions. The BC DNA was used as input material for PCR amplification followed by library preparation and deep sequencing. Primers were designed to amplify a 302bp region of the rat *Th* promoter, including exon 1 (−139 to +164 bp from the *Th* TSS) (Supplementary Table 1). Amplification was carried out on a C1000 Thermal Cycler (Bio-Rad, Hercules, CA) with 20 ng of BC DNA per reaction. The amplification conditions were: 40 cycles of 94°C for 30sec, 55°C for 30sec and 72°C for 1min sequencing libraries were prepared using the NETflex DNA Sequencing Kit (BIOO Scientific, Austin, TX) according to manufacturer’s instructions. The libraries were evaluated using an Agilent 2100 Bioanalyzer (Agilent Technologies, Palo Alto, CA) and were normalized to 2nM with 10 mM Tris-HCl. The libraries were then pooled and sequenced on a MiSeq (Illumina, Inc. San Diego, CA) by the Molecular & Cellular Biology Core, (ONPRC, Beaverton, OR) to generate 250base, paired-end reads. The reads were trimmed using Trim Galore^59^ and aligned to the reference genome using Bismark^60^. Alignment data was converted to CpG methylation rate using the Bismark methylation extractor and custom scripts.

### Immunohistochemistry

To assess the expression of proteins related to maternal behavior, immunoreactivity of TH was measured in EDC mixture exposed and non-exposed females from the F1 generation at P21. TH-ir was measured in the substantia nigra (SN), the ventral tegmental area (VTA) and the median preoptic area (mPOA). Females were anesthetized with pentobarbital sodium (50 mg kg^−1^, i.p.) and sequentially perfused with PBS and 4% paraformaldehyde. Brains were subsequently removed and post-fixed in 4% paraformaldehyde at 4 °C overnight. Therefore, coronal sections (30 μm) were cut on a vibratome and used for immunohistofluorescence.

For immunohistochemistry, sections were incubated in PBS and blocked with 10% donkey serum for 1 h at room temperature, and incubated overnight with the primary antibody anti-TH (Th-mouse, 22941, ImmunoStar, Hudson, USA, 1:1000) for 24h at room temperature or 4 °C. Thereafter, the corresponding fluorophore-conjugated secondary antibodies were incubated for 2 h at room temperature. The high-resolution Zeiss microscope LSM880 implemented with the fast Airyscan detector was used for visualization. Three slides per animal per region of interest (ventral midbrain sections containing the SN and VTA or median preoptic area sections) were used for quantification. An observer blind to condition outlined each region of interest for nuclei-specific analysis. Region area and number of immunoreative cells were automatically quantified using Imaris 9.3. Statistics were performed on the total cell count per animal per region across slice sections.

### Massively parallel RNA sequencing

Poly-A mRNA was purified from 1 microgram total RNA using NETFLEX poly-A beads (Perking-Elmer, Austin, TX) followed by library preparation using the NETFLEX Rapid Directional RNA-seq Kit 2.0 (Perking-Elmer). In short, after fragmentation with divalent cations and heat, mRNA was used as template for reverse transcription using random hexamer primers. cDNAs were then blunted and 3’ end A-tailed to facilitate adaptor ligation. Six-base pair Illumina adaptors were ligated followed by 12 rounds of PCR amplification. Free dNTPs were removed using AMPure XP beads (BeckmanCoulter, Brea, CA). Distribution of DNA sizes in the library was confirmed by Bioanalyzer analysis (Agilent, Santa Clara, CA). Library titer was determined by real-time PCR (Kapa Biosystems, Wilmington, MA) on a Quant Studio 12K Flex Real Time PCR System (ThermoFisher, Waltham, MA). Four samples were sequenced per lane on a HiSeq 4000 (Illumina). Sequencing was done using a single-read 100-cycle protocol. The resulting base call files (.bcl) were converted to standard fastq formatted sequence files using Bcl2Fastq (Illumina). Sequencing quality was assessed using FastQC (Babraham Bioinformatics, Cambridge, UK). The RNAseq procedure was carried out by the Genomics & Cell Characterization Core Facility at the University of Oregon. To determine the differential gene expression values we used the gene-level edgeR analysis package. We performed an initial trimming and adapter removal pass using Trimmomatic. After this reads were aligned to the rn6 build of the rat genome with Bowtie2/Tophat2, and assigned to gene-level genomic features with the Rsubread featureCounts package based on the Ensembl 83 annotation set. Differential expression between experimental groups was analyzed using the generalized linear modeling approaches implemented in edgeR. Lists of differentially expressed genes/transcripts were identified based on significance of pairwise comparison of experimental groups. Gene ontology and enrichment analysis were performed using the database for annotation, visualization and integrated discovery (DAVID).

### ChIP assay

To assess activatory and repressive histone modifications at specific gene promoters affected by EDC mixture exposure we performed ChIP assay using extracted chromatin from the hypothalamus of prepubertal rats at P21. As described previously^14,61^, ChIP procedure was carried out by crosslinking tissue in phosphate-buffered saline (PBS) containing a protease inhibitor cocktail (PI, 1 mM phenylmethylsulfonylfluoride, 7 μg ml−1 aprotinin, 0.7 μg ml−1 pepstatin A, and 0.5 μg ml−1 leupeptin), a phosphatase inhibitor cocktail (PhI, 1 mM β-glycerophosphate, 1 mM sodium pyrophosphate, and 1 mM sodium fluoride), and an HDAC inhibitor (20 mM sodium butyrate) at 4°C and 1% formaldehyde for 10 min at room temperature. After two washing steps in PBS, samples were lysed with 200 µl SDS buffer (0.5% SDS, 50 mM Tris-HCl, and 10 mM EDTA) containing protease, phosphatase, and HDAC inhibitors, and sonicated for 45 s to yield chromatin fragments of ∼500 base pairs (bp) using the microtip of a Fisher Scientific FB 705 sonicator. Size fragmentation was confirmed by agarose gel electrophoresis. The sonicated chromatin was clarified by centrifugation at 14 000 r.p. m. for 10 min at 4 °C, brought up to 1 ml in Chip Dilution Buffer (16.7 mM TrisHCl, pH 8.1, 150 mM NaCl, 1.2 mM EDTA, 1.1% Triton X-100, and 0.01% SDS) containing the PI and PhI cocktails, and the HDAC inhibitor described above. The samples were then stored at −80 °C for subsequent immunoprecipitation. For this step, chromatin was pre-cleared with Protein A/G beads (Dynabeads, Invitrogen) for 1 h at 4 °C. Twenty-five to 50 μl aliquots of chromatin were then incubated with 2–5 μg of the antibodies described in Supplementary Table 2. The complexes were incubated with 25 μl of protein A or G beads solution (Dynabeads) at 4 °C overnight with mild agitation. The next day the beads were washed first with 0.5 ml low-salt wash buffer (20 mM Tris-HCl, pH 8.1, 150 mM NaCl, 2 mM EDTA, 1% Triton X-100, and 0.1% SDS), followed by high-salt wash buffer (20 mM Tris-HCl, pH 8.1, 500 mM NaCl, 2 mM EDTA, 1% Triton X-100, and 0.1% SDS), LiCl buffer (10 mM Tris-HCl, pH 8.1, 250 M LiCl, 1% Nonidet P-40, 1% sodium deoxycholate, and 1 mM EDTA), and finally with TE buffer (10 mM Tris-HCl, pH 8.0, and 1 mM EDTA). Thereafter, the immunocomplexes were eluted with 100 μl of 0.1 M NaHCO3 and 1% SDS at 65 °C for 45 min. To reverse the crosslinking reaction, we added 4 μl of 5 M NaCl and incubated the samples at 95 °C for 30 min. We recovered the DNA using ChIP DNA Clean & Concentrator columns (Zymo Research, Irvine, CA), and stored the resulting material at −80 °C before qPCR analysis. All the chemicals mentioned above were purchased from Sigma-Aldrich.

### qPCR detection of chromatin immunoprecipitated DNA

Genomic regions of interest were amplified by qPCR. Primer sequences, accession numbers of the genes analyzed as well as the chromosomal position of the 5′-flanking region amplified, using the position of the TSS as the reference point, are shown in Supplementary Table 1. PCR reactions were performed using 1 μl of each immunoprecipitate (IP) or input samples, primer mix (1 µM each primer), and SYBR Green Power Up Master Mix™ (Thermo Fisher) in a final volume of 10μl. Input samples consisted of 10% of the chromatin volume used for immunoprecipitation. The thermocycling conditions used were as follows: 95°C for 5 min, followed by 40 cycles of 15sec at 95°C and 60sec at 60°C. Data are expressed as % of IP signal/input signal.

### Statistics

All statistical analyses were performed using Prism 7.0 software (GraphPad, San Diego, CA). Data was subjected to a normality and an equal variance test and parametric test were used when conditions were accomplished. Parametric test used were one-way or two-way ANOVA followed by Student–Newman– Keuls or Sidak’s for multiple comparisons, respectively; or the Student’s t-test to compare two groups. When comparing percentages, groups were subjected to an arcsine transformation before statistical analysis to convert the values from a binomial to a normal distribution. Data that did not accomplish normality followed a Mann-Whitney test. When making multiple comparisons α was adjusted by using the Bonferroni correction. The investigator was group blinded in all physiological and molecular determination. Samples size, reported in figure captions, were estimated based in previous studies or based in the calculation of an adequate statistical power. Effect sizes were calculated according Cohen’s delta formula (Supplementary Table 3-6).The level of statistical significance was a P value <0.05.

## ACKNOWLEDGEMENTS

We will be forever grateful for Professor Jean-Pierre Bourguignon’s mentorship whose guidance and sharp scientific mind will be greatly missed. His ideas and insights deeply improved the quality of this project from its conception. We are indebted to Professor P. Delvenne for assistance with Papanicolaou staining and to Dr. V. D. Ramirez (Urbana, IL) for providing the CR11-B81 anti-GnRH antiserum. This work was supported by grants from the NIH (1R01HD084542) to A.L and 8P51OD011092 for the operation of the Oregon National Primate Research Center as well as the FRS-FNRS (“Fonds Nationale de la Recherche Scientifique” (CDR-J013019F-33661942) and the Belgian Society for Pediatric Endocrinology and Diabetology.

## AUTHOR CONTRIBUTIONS

D.L.R. contributed to the study design and writing of the manuscript, conducted the physiological, molecular and epigenetic experiments, the statistical analysis, evaluated and discussed the data; C.F.A helped to perform RNA-seq and ChIP experiments; V.D., M.M, M.C, E.S and A.G were involved in physiological experiments and data analysis. A.L and A.S.P supervised the study design, analyzed the data and wrote the manuscript, which was revised by the rest of the authors. All the authors take full responsibility for the work.

