## Supplemental material for "Multi- and transgenerational disruption of maternal behavior and female puberty by Endocrine Disrupting Chemical (EDC) mixture exposure"

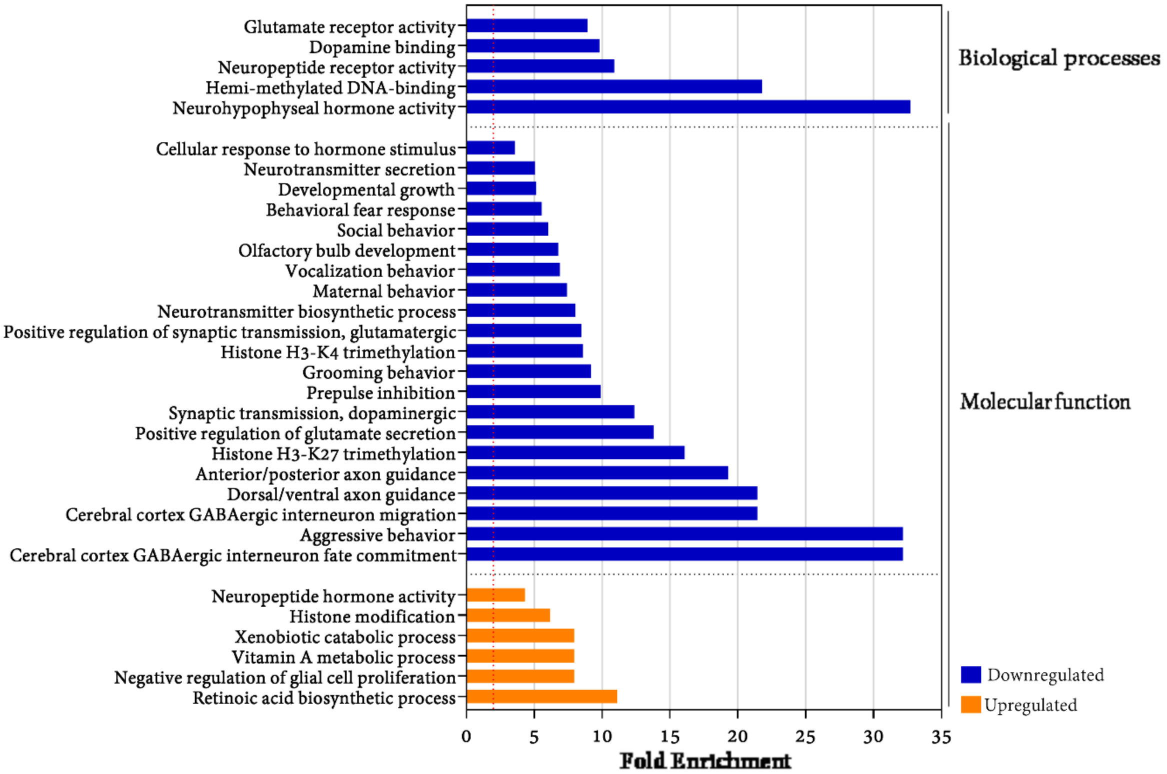


**Supplementary Figure 1.** Fold enrichment of gene ontology (GO) annotations using David pathway analysis across compared groups (CTL vs EDC) in the MBH of F3 generation females at P21. The gene enrichment analysis grouped the differentially expressed genes using GO annotations data. We selected enriched GO annotations using 2-fold enrichment criteria as a threshold and identifying annotations that were involved in brain and behavioral processes. Those annotations were then categorized in upregulated (orange) or downregulated (blue) annotations.

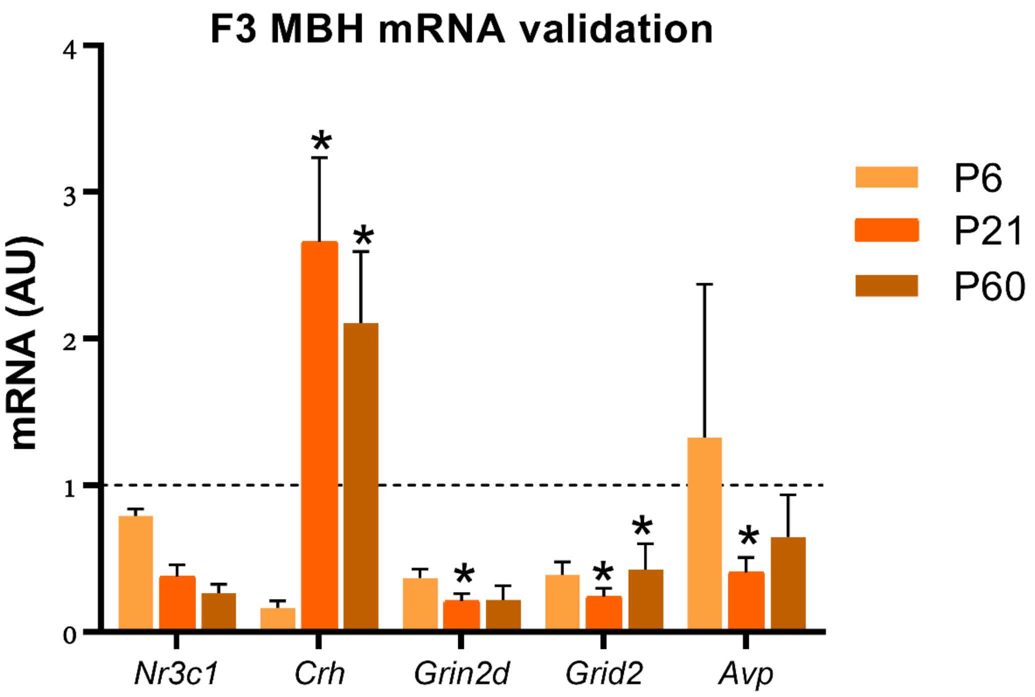


**Supplementary Figure 2.** *Nr3c1*, *Crh*, *Grin2d*, *Grid2* and *Avp* mRNA expression in the female rat ancestrally (F3 generation) exposed to an EDC mixture or vehicle in the MBH of infant (P6), prepubertal (P1) and adult (P60) female rats as determined by qPCR (n=6/group). AU = arbitrary units. RNA expression data were normalized by dividing each individual value by the average of the control group at every time point. Bars represent mean ± s.e.m. (*P < 0.05, **P < 0.01, ***P < 0.001 vs. CTL, Student's t-test).

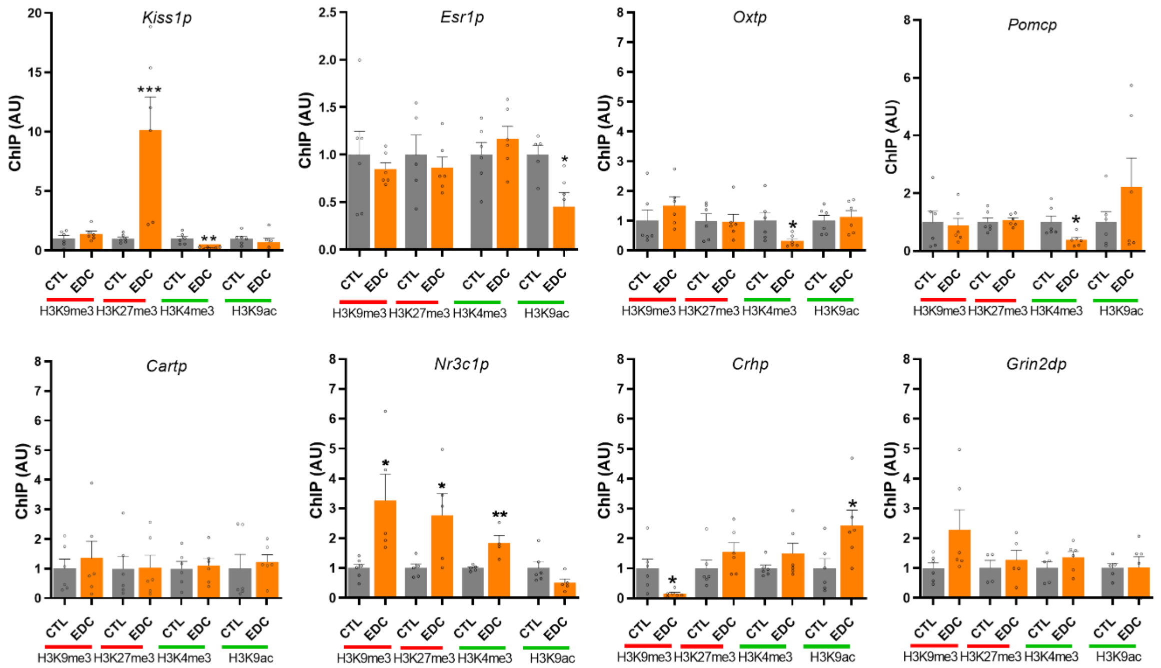


**Supplementary figure 3.** Abundance of the TrxG-dependent activating marks H3K4me3 and H3K9ac and the PcG-dependent repressive mark H3K27me3 and H3K9me3 at the *Kiss1*, *Esr1*, *Oxt*, *Pomc*, *Cart*, *Nr3c1*, *Crh* and *Grin2d* promoter in the prepubertal MBH of females EDC and control from the F3 generation, as measured by ChIP (n=6/group). Bars represent mean ± s.e.m. (*P < 0.05, **P < 0.01, ***P < 0.001 vs. CTL, Student's t-test).

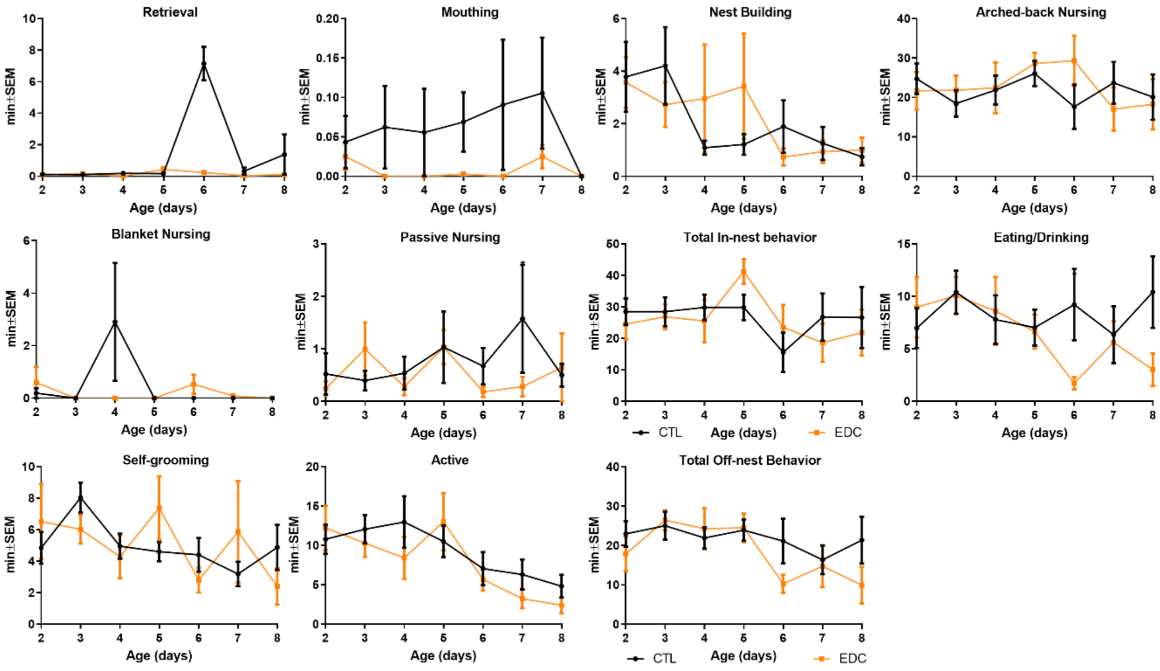


**Supplementary figure 4.** Maternal behavior displayed by female rats directly exposed to a mixture of EDC from the F0 generation. Data shows time spent by dams displaying in-nest behavior (retrieval, mouthing, nest building, Arched-back, blancked and passive nursing; and total time spent in-nest) or off-nest behaviors (eating/drinking, self-grooming, being active and total time off-nest) from P2 to P8. Plotted lines represent average of time ± s.e.m.

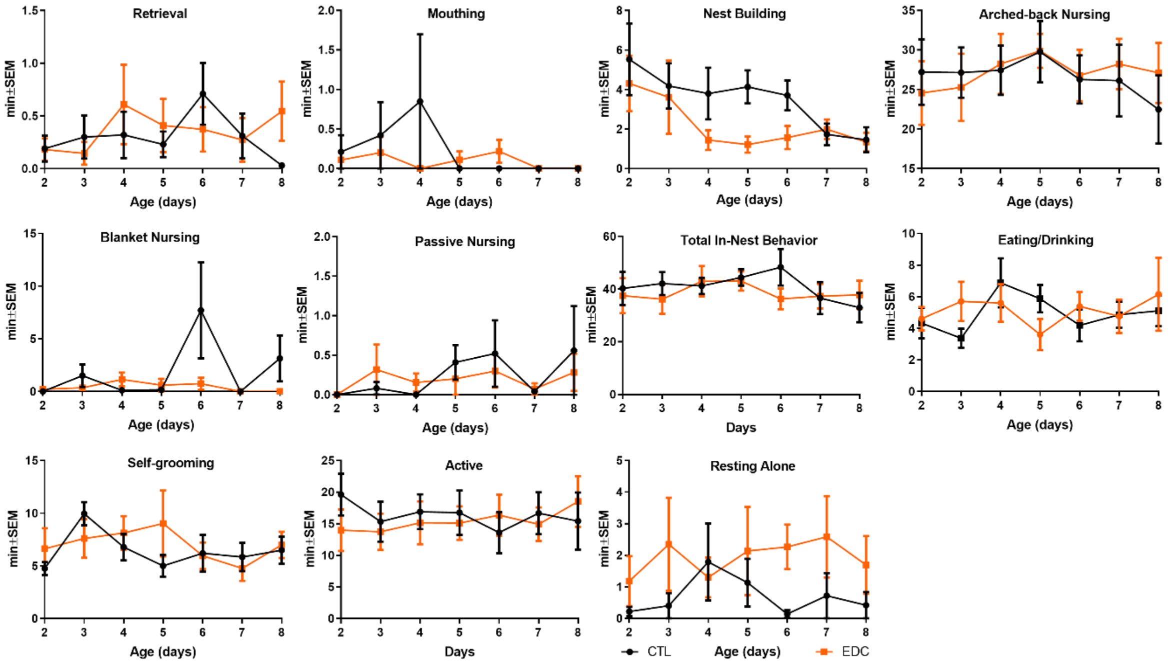


**Supplementary figure 5.** Maternal behavior displayed by female rats directly exposed to a mixture of EDC from the F1 generation. Data shows time spent by dams displaying in-nest behavior (retrieval, mouthing, nest building, Arched-back, blancked and passive nursing; and total time spent in-nest) or off-nest behaviors (eating/drinking, self-grooming, being active and total time off-nest) from P2 to P8. Plotted lines represent average of time ± s.e.m.

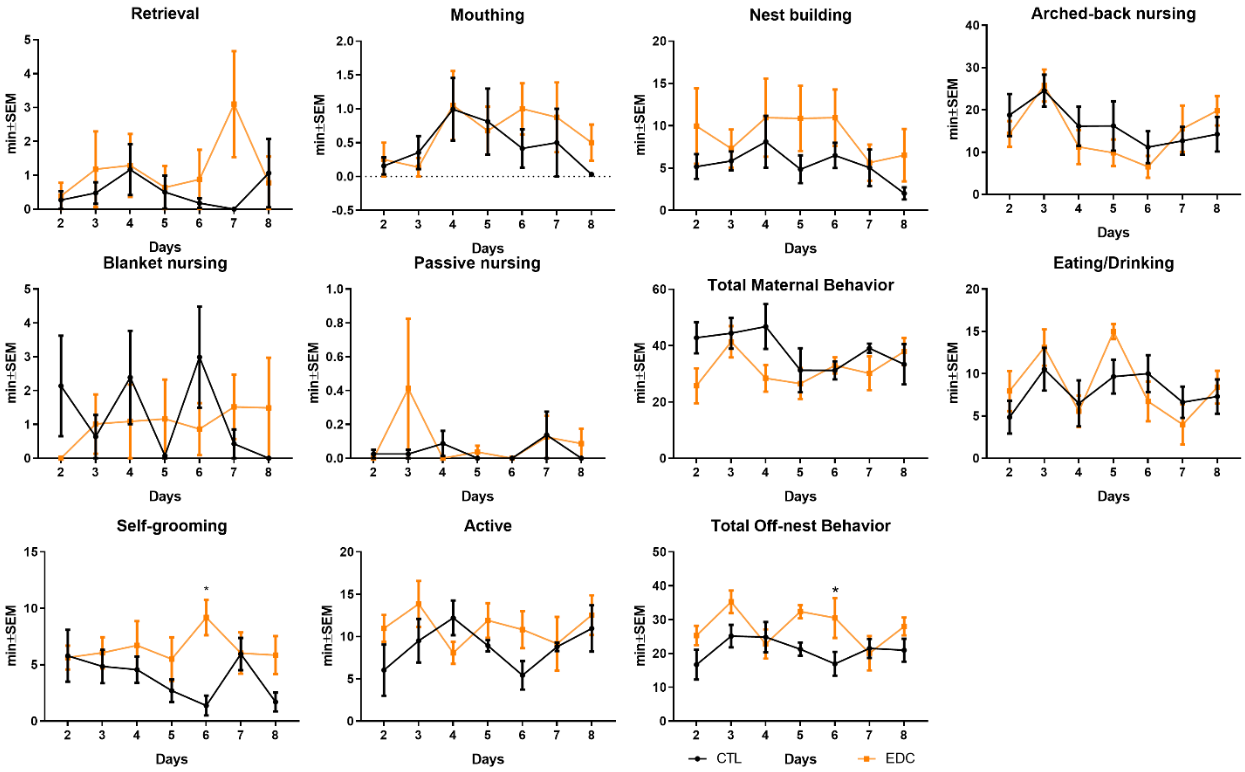


**Supplementary figure 6.** Maternal behavior displayed by female rats directly exposed to a mixture of EDC from the F2 generation. Data shows time spent by dams displaying in-nest behavior (retrieval, mouthing, nest building, Arched-back, blancked and passive nursing; and total time spent in-nest) or off-nest behaviors (eating/drinking, self-grooming, being active and total time off-nest) from P2 to P8. Plotted lines represent average of time ± s.e.m.

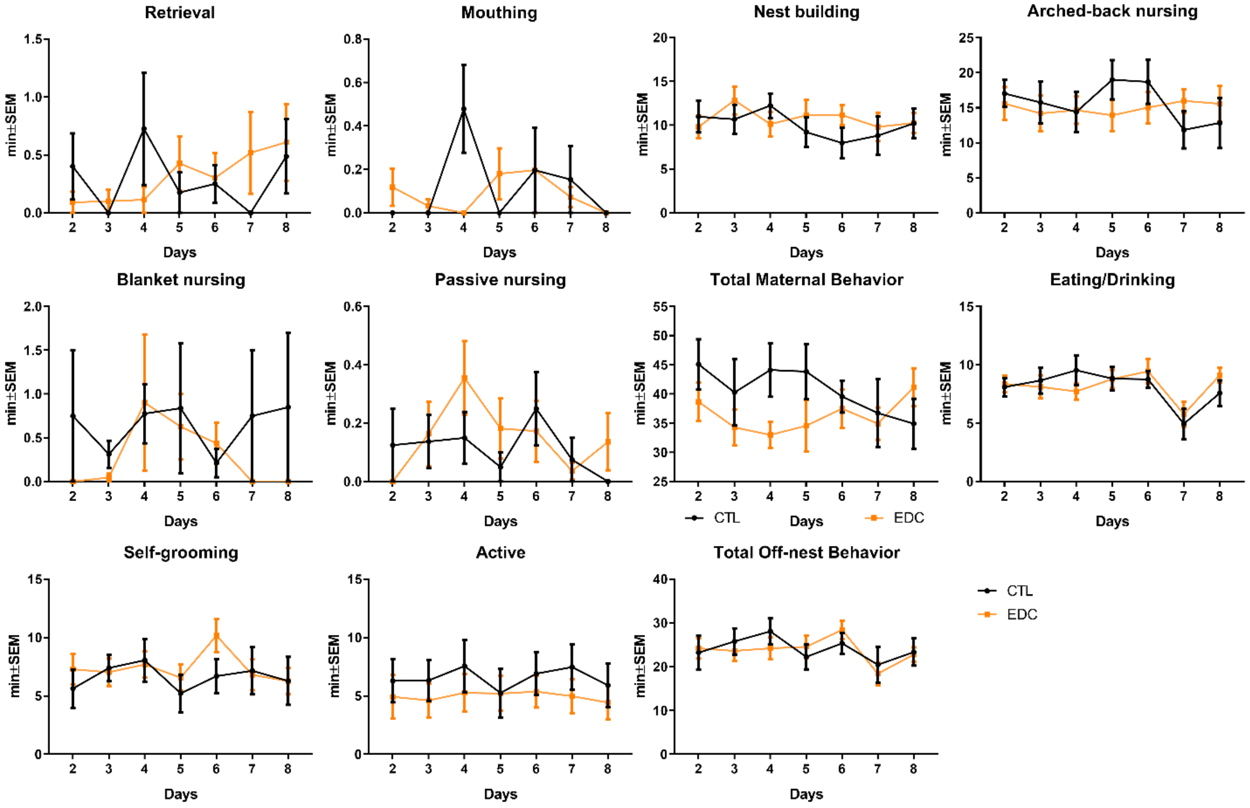


**Supplementary figure 7.** Maternal behavior displayed by female rats directly exposed to a mixture of EDC from the F3 generation. Data shows time spent by dams displaying in-nest behavior (retrieval, mouthing, nest building, Arched-back, blancked and passive nursing; and total time spent in-nest) or off-nest behaviors (eating/drinking, self-grooming, being active and total time off-nest) from P2 to P8. Plotted lines represent average of time ± s.e.m.

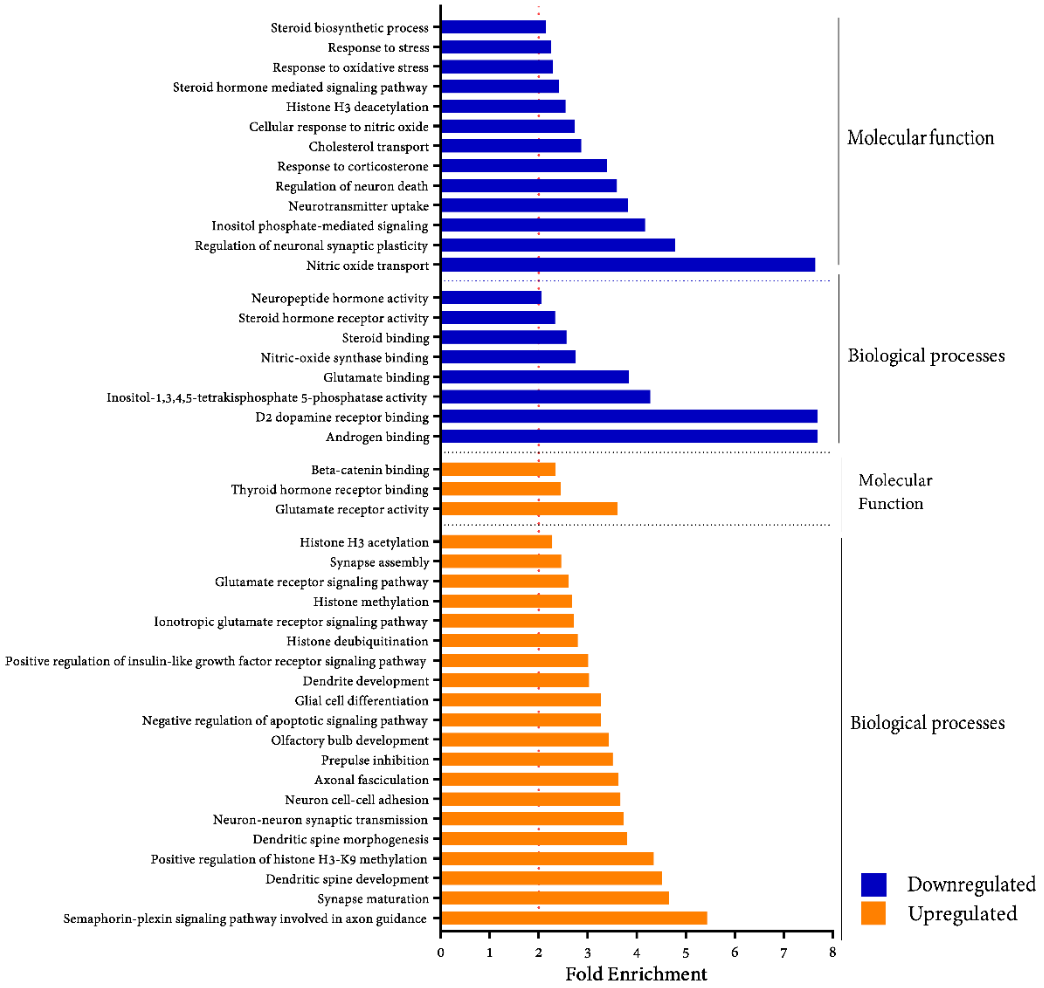


**Supplemental figure 8.** Fold enrichment of gene ontology (GO) annotations using David pathway analysis across compared groups (CTL vs EDC) in the MBH of F1 generation females at P21 . The gene enrichment analysis grouped the differentially expressed genes using GO annotations data. We selected enriched GO annotations using 2-fold enrichment criteria as a threshold and identifying annotations that were involved in brain and behavioral processes. Those annotations were then categorized in upregulated (orange) or downregulated (blue) annotations.

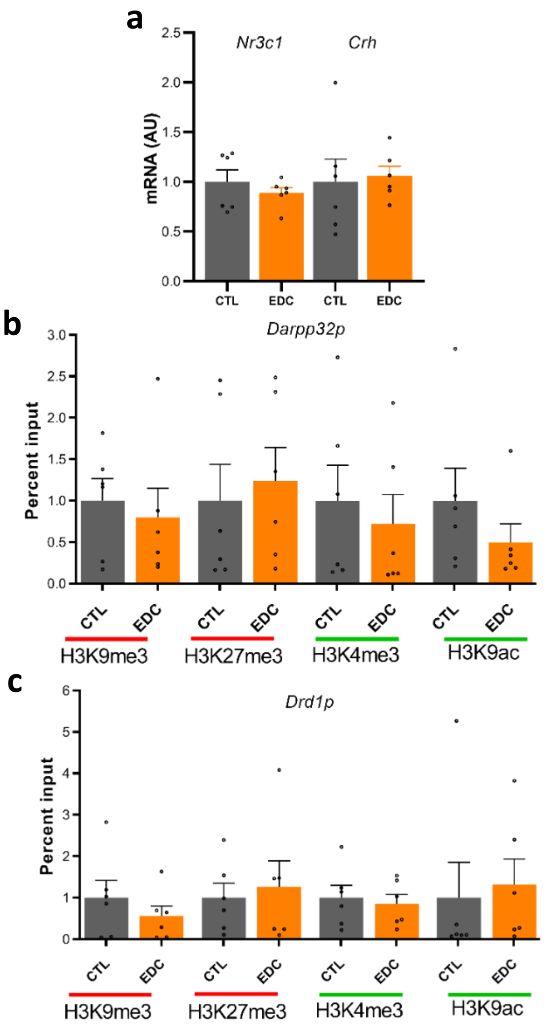


**Supplemental figure 9.** *Nr3c1* and *Crh* mRNA expression and *Darpp32* and *Dard1* promoter chromatin state in the female rat *in utero* and lactationally (F1 generation) exposed to an EDC mixture or vehicle. (**a**) Expression of *Nr3c1* and *Crh* mRNA in the MBH prepubertal (P21) female rats as determined by qPCR (n=6/group). AU = arbitrary units. RNA expression data were normalized by dividing each individual value by the average of the control group at every time point. (**b-c**) Abundance of the TrxG-dependent activating marks H3K4me3 and H3K9ac and the PcG-dependent repressive mark H3K27me3 and H3K9me3 at the *Darpp32* and *Drd1* promoter in the prepubertal MBH of females perinatally exposed to a mixture of EDC (F1 generation), as measured by ChIP (n=6/group). Bars represent mean ± s.e.m.

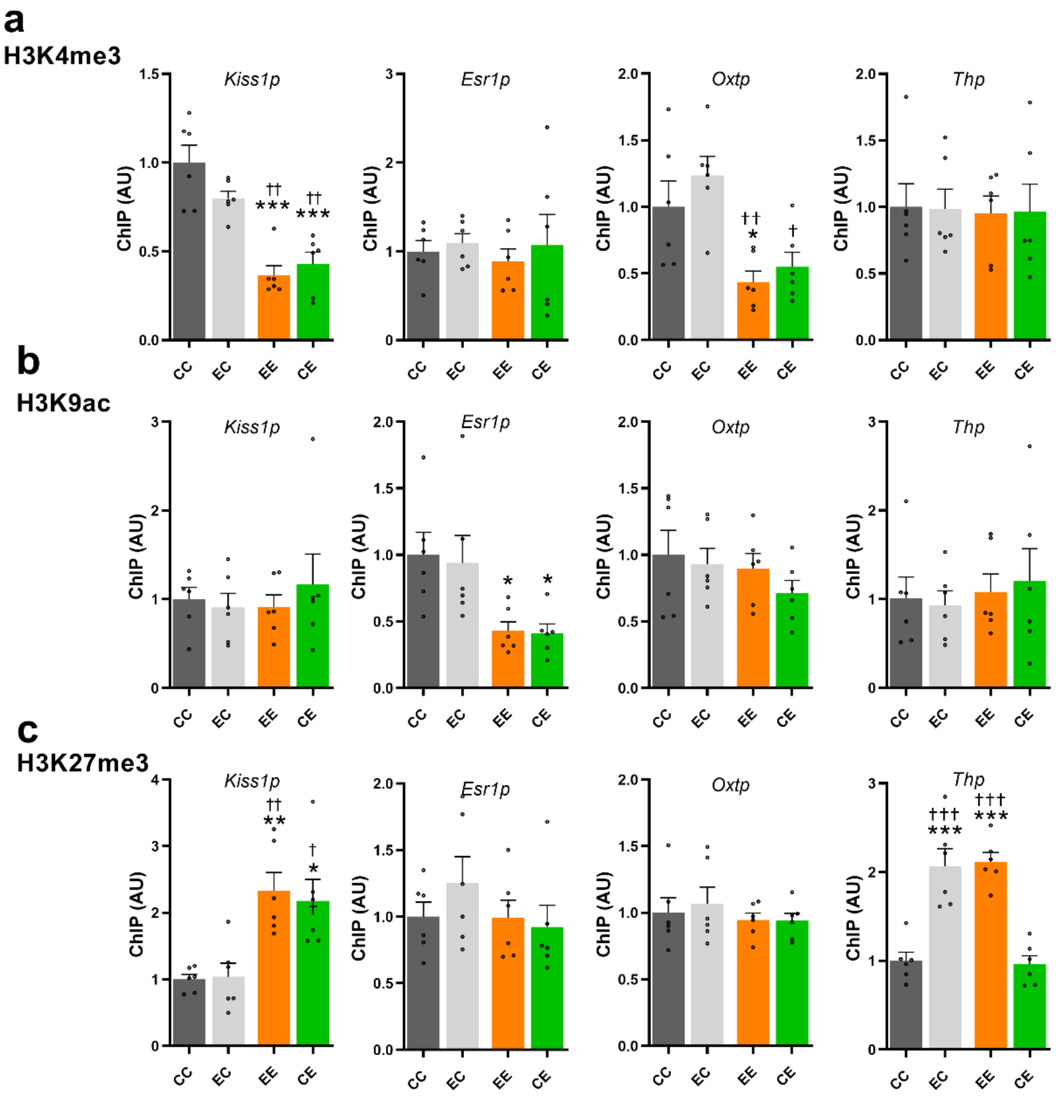


**Supplemental figure 10.** Abundance of the TrxG-dependent activating marks H3K4me3 (a) and H3K9ac (b) and the PcG-dependent repressive mark H3K27me3 (c) at the *Kiss1*, *Esr1*, *Oxt* and *Th* promoter in the prepubertal (P21) MBH of cross-fostered germ-cell EDC exposed pups or control (F2 generation) raised by either *in utero* EDC exposed dams or control, as measured by ChIP (n=6/group). CC= control pup raised by control dam; EC: control pup raised by *in utero* EDC exposed dam; EE= germ-cell EDC exposed pup raised raised by *in utero* EDC exposed dam; CE= germ-cell EDC exposed pup raised by control dam. Bars represent mean ± s.e.m. (*P < 0.05, **P < 0.01 vs. CC, †P < 0.05 vs. CE, one-way ANOVA).

Supplemental Table 1. Primer Sequence

| Gene | | Primer | Accession number | Amplicon size | Use |
|---|---|---|---|---|---|
| *Kiss1* | F<br>R | *TGGTGAACCCTGAACCCACAGGC*<br>*CGGCGGGCATGGCGATGTT* | NM_181692.1 | 136 | qPCR |
| *Esr1* | F<br>R | *CGCTCTGCCTTGATCACACA*<br>*GCCGAGGTACAGATTGGCTT* | NM_012689.1 | 188 | qPCR |
| *Oxt* | F<br>R | *GCTGCCAGGAGGAGAACTAC*<br>*ATCATCACAAAGCGGGCTCA* | NM_012996.3 | 175 | qPCR |
| *Pomc* | F<br>R | *CTTTCCGCGACAGAGCCT*<br>*CCAGCTCCACACGTCTATGG* | NM_139326.2 | 113 | qPCR |
| *Th* | F<br>R | *CCTTCCAGTACAAGCACGGT*<br>*TGGGTAGCATAGAGGCCCTT* | NM_012740.3 | 109 | qPCR |
| *Cart* | F<br>R | *GCGCTGTGTTGCAGATTGAA*<br>*CGTCACACATGGGGACTTGG* | NM_017110.1 | 105 | qPCR |
| *Nr3c1* | F<br>R | *GGTGATTGAACCCGAGGTGT*<br>*TTTCTGAAGCCTGGTATCGCC* | NM_012576.2 | 147 | qPCR |
| *Crh* | F<br>R | *CAAGGGAGGAGAAGAGAGCG*<br>*AAGAAATTCAACGGCTGCGG* | NM_031019.1 | 160 | qPCR |
| *Grin2d* | F<br>R | *AGCTCTGCGACCTGCTGT*<br>*CCAAGCTGCAGGAAGGTGGA* | NM_022797.1 | 190 | qPCR |
| *Dnm1* | F<br>R | *TCTACAAGGATTACCGGCAGC*<br>*GCTTTCTCCTTGTCCCCAACA* | NM_080689.4 | 121 | qPCR |
| *Drd1* | F<br>R | *CCACTCTCCTGGGCAATACC*<br>*AAAAGGACCCAAAGGGCCAA* | XM_006253600.3 | 180 | qPCR |
| *Darpp32* | F<br>R | *CCCAAGGACCGCAAGAAGAT*<br>*CTCCTGAGGTTCTCTGGTGC* | NM_138521,1 | 172 | qPCR |
| *Grid2* | F<br>R | *GTCCCATCGAAAGAGGATGACA*<br>*ACTGTTTATGGGGGCTGTCG* | NM_024379.1 | 97 | qPCR |
| *Avp* | F<br>R | *AGCGATGAGAGCTGCGTG*<br>*CTGTACCAGCCTAAGCAGCA* | NM_016992.2 | 129 | qPCR |
| *Kiss1* | F<br>R | *TCGGGCAGCCAGATAGAGGAAGC*<br>*TTGAGGGCCGAGGGAGAAGAG* | NM_181692.1 | 91 | ChIP |
| *Esr1* | F<br>R | *GTCCCTCAGCAGCCAGCCAGTCT*<br>*CTCTCGGGAAGCAGCCAGTAGG* | NM_012689.1 | 127 | ChIP |
| *Oxt* | F<br>R | *TGTAGCTTAGGCCTCCCCTT*<br>*CATGACTGGTCACAGCAGGT* | NM_012996.3 | 159 | ChIP |
| *Pomc* | F<br>R | *GCTAAGCCTCTGTCCAGTCC*<br>*GTTAGCACAGACCCGCTGAA* | NM_139326.2 | 103 | ChIP |
| *Th* | F<br>R | *CCGACTGGGGCAGTGAATAG*<br>*TAACCAAACCAGGGCACACA* | NM_012740.3 | 198 | ChIP |
| *Cart* | F<br>R | *TTCCATTTCATGGGCCCTCC*<br>*GGCTGGAGCACAGAGAACAA* | NM_017110.1 | 139 | ChIP |
| *Nr3c1* | F<br>R | *AAGGGTTAGAAGGAATTTGGGGA*<br>*TGACGTGCCAGAGCCAATTA* | NM_012576.2 | 180 | ChIP |
| *Crh* | F<br>R | *ACGCAATCGAGCTGTCAAGA*<br>*CAGAGCCCGGAGTGAGATTT* | NM_031019.1 | 96 | ChIP |
| *Grin2d* | F<br>R | *TCTGGTTCTGTTCCTGGGTTTTTG*<br>*TGGGGTCAGGGAAGATACAGAGGT* | NM_022797.1 | 121 | ChIP |
| *Th − BS* | F<br><br>R | *TCGTCGGCAGCGTCAGATGTGTATA*<br>*AGAGACAGGTTTTTTTTAGGTATAGTAGG*<br>*GTCTCGTGGGCTCGGAGATGTGTATAA*<br>*GAGACAGtatttatttataggtacaaaag* | NM_012740.3 | 302 | BS-seq |

Supplemental Table 2. List of primary antibodies

| Target | Host | Source | Catalog # | Use |
|---|---|---|---|---|
| Th | Mouse | ImmunoStar | 22941 | IHC |
| H3K27me3 | Rabbit | Active Motif | 39155 | ChIP |
| H3K9ac | Rabbit | Active Motif | 39917 | ChIP |
| H3K4me3 | Rabbit | Active Motif | 39159 | ChIP |
| H3K9me3 | Mouse | Active Motif | 61013 | ChIP |
| $\beta$-Galatocidase | Rabbit | Cortex Biochem | CR7001RP2 | ChIP |
| $\beta$-Galatocidase | Mouse | ICN Biomedical | 55976 | ChIP |

Supplemental Table 3. Report of descriptive and statistical data.

| Measure | F | N | Average | | SD | | *p-value* | Effect | Figure |
|---|---|---|---|---|---|---|---|---|---|
| | | | CTL | EDC | CTL | EDC | | Size | |
| Vaginal Opening | F1 | 51/56 | 33,92 | 33,98 | 1,71 | 2,94 | | 0,02 | 2a |
| | F2 | 50/52 | 34,73 | 37,93 | 1,33 | 1,58 | 0,000 | 2,19 | 2b |
| | F3 | 15/24 | 34,29 | 38,20 | 1,92 | 1,32 | 0,000 | | 2c |
| | F4 | 47/64 | 31,36 | 34,50 | 1,36 | 1,40 | 0,000 | 2,27 | 2d |
| GnRH IP | F1 | 4 | 44,38 | 43,75 | 1,25 | 1,44 | | | 2a |
| | F3 | | 42,03 | 44,06 | 0,94 | 1,08 | 0,030 | 2,01 | 2c |
| Regular cycle | F1 | 20 | 88,75 | 89,38 | 27,77 | 26,06 | | 0,02 | 3a left |
| Proestrus | | | 23,93 | 23,31 | 4,06 | 5,18 | | -0,13 | |
| Estrus | | | 26,58 | 28,78 | 6,16 | 12,48 | | 0,22 | |
| Diestrus | | | 49,49 | 47,91 | 5,70 | 8,47 | | -0,22 | |
| Regular cycle | F2 | 15 | 89,17 | 60,83 | 24,49 | 42,75 | 0,034 | -0,81 | 3b left |
| Proestrus | | | 23,75 | 22,50 | 3,30 | 4,13 | | -0,33 | |
| Estrus | | | 26,04 | 40,42 | 4,96 | 17,34 | 0,001 | 1,13 | |
| Diestrus | | | 50,21 | 37,08 | 5,60 | 15,12 | 0,002 | -1,15 | |
| Regular cycle | F3 | 15/14 | 90,83 | 49,11 | 22,89 | 45,85 | 0,004 | -1,15 | 3c left |
| Proestrus | | | 24,08 | 17,73 | 5,24 | 8,87 | 0,048 | -0,87 | |
| Estrus | | | 26,29 | 37,79 | 1,70 | 12,59 | 0,000 | 1,28 | |
| Diestrus | | | 49,65 | 44,49 | 4,57 | 6,81 | 0,048 | -0,89 | |
| Folliculogenesis | | | | | | | | | 3a middle |
| Primordial | F1 | 10/9 | 17,83 | 12,40 | 11,52 | 8,13 | | -0,55 | |
| Primary | | | 6,28 | 6,68 | 4,41 | 5,95 | | 0,08 | |
| Secondary | | | 3,55 | 2,71 | 1,84 | 1,57 | | -0,49 | |
| Antral | | | 1,23 | 1,00 | 0,98 | 0,89 | | -0,25 | |
| Atretic | | | 8,42 | 16,25 | 3,87 | 11,70 | | 0,90 | |
| Corpora lutea | | | 0,02 | 0,05 | 0,03 | 0,05 | | 0,61 | |
| Cysts | | | 1,11 | 0,66 | 0,80 | 0,36 | | -0,73 | |
| Folliculogenesis | | | | | | | | | 3b middle |
| Primordial | F2 | 10/9 | 21,70 | 9,09 | 16,52 | 7,62 | 0,038 | -0,98 | |
| Primary | | | 9,25 | 9,08 | 6,73 | 5,61 | | -0,03 | |
| Secondary | | | 8,54 | 7,16 | 6,50 | 4,14 | | -0,25 | |
| Antral | | | 2,55 | 0,74 | 1,16 | 0,88 | 0,045 | -1,76 | |
| Atretic | | | 18,08 | 33,62 | 6,79 | 25,12 | 0,014 | 0,84 | |
| Corpora lutea | | | 0,01 | 0,12 | 0,03 | 0,16 | | 0,97 | |
| Cysts | | | 0,63 | 1,13 | 1,06 | 1,36 | 0,470 | 0,41 | |
| Folliculogenesis | | | | | | | | | 3c middle |
| Primordial | F3 | 10/9 | 25,31 | 9,96 | 12,37 | 5,59 | 0,033 | -1,60 | |
| Primary | | | 7,00 | 8,25 | 5,37 | 8,31 | | 0,18 | |
| Secondary | | | 2,49 | 3,52 | 1,98 | 2,21 | | 0,49 | |
| Antral | | | 1,20 | 0,29 | 0,85 | 0,37 | 0,048 | -1,38 | |
| Atretic | | | 10,64 | 13,70 | 10,06 | 8,08 | 0,002 | 0,34 | |
| Corpora lutea | | | 0,04 | 0,07 | 0,08 | 0,12 | | 0,27 | |
| Cysts | | | 1,13 | 1,02 | 0,60 | 1,31 | | -0,10 | |
| Ovarian Weight | F1 | 14 | 47,69 | 40,33 | 7,34 | 9,94 | 0,035 | -0,84 | 3a right |
| | F2 | 16 | 41,44 | 35,54 | 8,85 | 6,61 | 0,041 | -0,76 | 3b right |
| | F3 | 10/9 | 65,24 | 44,67 | 10,90 | 7,96 | 0,000 | -2,16 | 3c right |

**Notes**: Effect size was calculated using Cohen's *d*. F= generation. N=sample size. SD: standard deviation. IP: interpulse interval.

Supplemental Table 4. Report of descriptive and statistical data.

| **Measure** | **F** | **N** | **Average** | | **SD** | | ***p-value*** | **Effect Size** | **Figure** |
|---|---|---|---|---|---|---|---|---|---|
| | | | CTL | EDC | CTL | EDC | | | |
| *Kiss1* P6 | F3 | 6 | 1 | 0,39 | 0,13 | 0,23 | | -3,23 | 4a |
| *Kiss1* P21 | | | 1 | 0,51 | 0,34 | 0,29 | 0,034 | -1,56 | |
| *Kiss1* P70 | | | 1 | 0,38 | 0,07 | 0,34 | 0,026 | -2,52 | |
| *Esr1* P6 | | | 1 | 0,85 | 0,15 | 0,24 | | -0,76 | |
| *Esr1* P21 | | | 1 | 0,32 | 0,50 | 0,32 | 0,019 | -1,61 | |
| *Esr1* P70 | | | 1 | 0,21 | 0,07 | 0,16 | 0,013 | -6,49 | |
| *Oxt* P6 | | | 1 | 0,22 | 0,15 | 0,16 | | -4,95 | |
| *Oxt* P21 | | | 1 | 0,23 | 1,26 | 0,19 | 0,018 | -0,86 | |
| *Oxt* P70 | | | 1 | 0,36 | 0,09 | 0,33 | 0,043 | -2,64 | |
| *Cart* P6 | | | 1 | 0,77 | 0,10 | 0,78 | | -0,41 | |
| *Cart* P21 | | | 1 | 3,24 | 0,32 | 1,66 | 0,013 | 1,88 | |
| *Cart* P70 | | | 1 | 0,87 | 0,16 | 0,43 | | -0,41 | |
| *Pomc* P6 | | | 1 | 0,22 | 0,15 | 0,15 | | -5,24 | |
| *Pomc* P21 | | | 1 | 2,60 | 0,15 | 1,38 | 0,023 | 1,63 | |
| *Pomc* P70 | | | 1 | 1,00 | 0,17 | 0,75 | | 0,01 | |
| *Kiss1*p H3K27me3 | | | 1 | 10,15 | 0,34 | 6,78 | 0,008 | 1,91 | 4b |
| *Kiss1*p H3K4me3 | | | 1 | 0,22 | 0,46 | 0,09 | 0,002 | -2,38 | |
| *Esr1*p H3K9ac | | | 1 | 0,32 | 0,69 | 0,20 | 0,044 | -1,33 | |
| *Oxt*p H3K4me3 | | | 1 | 0,32 | 0,69 | 0,20 | 0,044 | -1,33 | |
| *Pomc*p H3K4me3 | | | 1 | 0,38 | 0,51 | 0,21 | 0,021 | -1,58 | |
| Licking | F0 | 15 | 6,52 | 5,14 | 2,64 | 3,07 | | -0,49 | 5a |
| | F1 | 10/11 | 9,18 | 5,84 | 3,27 | 3,59 | 0,038 | -0,97 | 5b |
| | F2 | 11 | 11,93 | 6,38 | 3,72 | 3,49 | 0,008 | -1,54 | 5c |
| | F3 | 11 | 13,79 | 9,71 | 2,40 | 2,79 | 0,004 | -1,57 | 5d |
| Resting alone | F0 | 15 | 0,57 | 0,24 | 1,09 | 0,35 | | -0,41 | 5a |
| | F1 | 10/11 | 0,68 | 1,93 | 0,93 | 1,07 | 0,011 | 1,24 | 5b |
| | F2 | 11 | 0,44 | 1,62 | 0,43 | 1,19 | 0,021 | 1,30 | 5c |
| | F3 | 11 | 2,85 | 3,18 | 3,20 | 3,14 | | 0,10 | 5d |
| *Th* P6 | F1 | 6 | 1 | 0,65 | 0,19 | 0,27 | | -1,51 | 6a |
| *Th* P21 | | | 1 | 0,18 | 0,64 | 0,10 | 0,031 | -1,79 | |
| *Th* P60 | | | 1 | 0,11 | 0,22 | 0,13 | 0,013 | -4,93 | |
| *Dnm1* P6 | | | 1 | 0,30 | 0,24 | 0,14 | | -3,58 | |
| *Dnm1* P21 | | | 1 | 0,33 | 0,10 | 0,26 | 0,039 | -3,45 | |
| *Dnm1* P60 | | | 1 | 1,15 | 0,29 | 0,66 | | 0,30 | |
| *Drd1* P6 | | | 1 | 1,22 | 0,22 | 0,33 | | 0,79 | |
| *Drd1* P21 | | | 1 | 3,41 | 0,24 | 1,70 | 0,016 | 1,99 | |
| *Drd1* P60 | | | 1 | 0,95 | 0,14 | 0,76 | | -0,09 | |
| *Darpp32* P6 | | | 1 | 0,75 | 0,17 | 0,29 | | -1,03 | |
| *Darpp32* P21 | | | 1 | 0,26 | 0,73 | 0,19 | | -1,39 | |
| *Darpp32* P60 | | | 1 | 0,19 | 0,30 | 0,09 | 0,002 | -3,64 | |
| *Th*p H3K27me3 | | | 1 | 3,12 | 0,52 | 2,10 | 0,019 | 1,39 | 6c |
| *Th*p H3K9me3 | | | 1 | 1,32 | 1,12 | 1,27 | | 0,26 | |
| *Th*p H3K4me3 | | | 1 | 1,19 | 0,27 | 0,77 | | 0,33 | |
| *Th*p H3K9ac | | | 1 | 1,21 | 0,64 | 0,67 | | 0,32 | |
| *Th*-ir SN | | | 92,7 | 87,64 | 21,05 | 21,15 | | -0,24 | 6e |
| *Th*-ir VTA | | | 125,3 | 116,20 | 26,47 | 24,22 | | -0,36 | |
| *Th*-ir mPoA | | | 3,98 | 2,53 | 0,61 | 0,36 | 0,001 | -2,90 | |
| *Th* P6 | F3 | | 1 | 0,40 | 0,89 | 0,37 | | -0,88 | 6f |
| *Th* P21 | | | 1 | 0,36 | 0,16 | 0,17 | 0,040 | -3,79 | |
| *Th* P60 | | | 1 | 0,46 | 0,91 | 0,44 | | -0,77 | |
| *Drd1* P6 | | | 1 | 0,36 | 0,58 | 0,23 | 0,486 | -1,47 | |
| *Drd1* P21 | | | 1 | 0,47 | 0,11 | 0,15 | 0,042 | -4,11 | |
| *Drd1* P60 | | | 1 | 0,15 | 0,17 | 0,25 | 0,001 | -3,93 | |
| *Th*p H3K27me3 | | | 1 | 2,16 | 0,47 | 1,62 | | 0,97 | 6h |
| *Th*p H3K9me3 | | | 1 | 2,17 | 0,38 | 1,13 | 0,037 | 1,38 | |
| *Th*p H3K4me3 | | | 1 | 0,90 | 0,36 | 0,43 | | -0,26 | |
| *Th*p H3K9ac | | | 1 | 1,04 | 0,33 | 0,51 | | 0,08 | |

**Notes**: Effect size was calculated using Cohen's *d*. F= generation. N=sample size. SD: standard deviation. qPCR and ChIP data were normalized to the control group.

Supplemental Table 5. Report of descriptive and statistical data.

| Measure | F | N | Group | Average | SD | Comparison | *p-value* | Effect Size | Figure |
|---|---|---|---|---|---|---|---|---|---|
| Vaginal opening | F2-C | 16/19 | CC | 34,67 | 1,56 | CC vs. CE | 0,003 | -1,29 | 7a |
| | | | CE | 37,05 | 2,09 | CC vs. EE | 0,003 | -1,46 | |
| | | | EE | 37,13 | 1,81 | CE vs. EC | 0,001 | 1,30 | |
| | | | EC | 34,72 | 1,45 | EE vs. EC | 0,001 | 1,47 | |
| Regular cycle | | 10 | CC | 85,00 | 18,34 | | | | 7b |
| | | | CE | 75,00 | 27,50 | | | | |
| | | | EE | 75,00 | 23,90 | | | | |
| | | | EC | 88,33 | 17,66 | | | | |
| Proestrus | | | CC | 21,25 | 3,65 | | | | |
| | | | CE | 19,58 | 6,53 | | | | |
| | | | EE | 22,08 | 4,83 | | | | |
| | | | EC | 22,92 | 2,95 | | | | |
| Estrus | | | CC | 23,33 | 3,51 | CC vs. EE | 0,008 | -1,13 | |
| | | | CE | 25,83 | 6,15 | | | | |
| | | | EE | 32,50 | 10,90 | | | | |
| | | | EC | 26,25 | 4,41 | | | | |
| Diestrus | | | CC | 55,42 | 6,23 | CC vs. EE | 0,002 | 1,30 | |
| | | | CE | 54,58 | 8,21 | CE vs. EE | 0,008 | 1,07 | |
| | | | EE | 45,42 | 8,88 | | | | |
| | | | EC | 50,83 | 4,30 | | | | |
| *Kiss1* | | 6 | CC | 1,00 | 0,19 | CC vs. CE | 0,001 | 2,94 | 7c |
| | | | CE | 0,46 | 0,18 | CC vs. EE | 0,012 | 2,03 | |
| | | | EE | 0,61 | 0,20 | CE vs. EC | 0,000 | -2,96 | |
| | | | EC | 1,05 | 0,22 | EE vs. EC | 0,004 | -2,13 | |
| *Esr1* | | | CC | 1,00 | 0,41 | CC vs. CE | 0,015 | 2,12 | 7d |
| | | | CE | 0,36 | 0,12 | CC vs. EE | 0,038 | 1,84 | |
| | | | EE | 0,44 | 0,13 | CE vs. EC | 0,006 | -2,03 | |
| | | | EC | 1,07 | 0,48 | EE vs. EC | 0,017 | -1,79 | |
| *Oxt* | | | CC | 1,00 | 0,22 | CC vs. CE | 0,003 | 2,68 | 7e |
| | | | CE | 0,42 | 0,22 | CC vs. EE | 0,005 | 2,87 | |
| | | | EE | 0,44 | 0,17 | CE vs. EC | 0,001 | -2,17 | |
| | | | EC | 1,06 | 0,36 | EE vs. EC | 0,002 | -2,22 | |
| *Th* | | | CC | 1,00 | 0,10 | CC vs. EE | 0,000 | 5,02 | 7f |
| | | | CE | 1,14 | 0,26 | CC vs. EC | 0,001 | 3,42 | |
| | | | EE | 0,37 | 0,15 | CE vs. EE | 0,000 | 3,68 | |
| | | | EC | 0,52 | 0,17 | CE vs. EC | 0,000 | 2,83 | |
| *Kiss1* | F3-C | | CC | 1,00 | 0,35 | CC vs. CE | 0,027 | 1,42 | 7c |
| | | | CE | 0,56 | 0,26 | CC vs. EE | 0,043 | 1,47 | |
| | | | EE | 0,59 | 0,18 | CE vs. EC | 0,007 | -2,53 | |
| | | | EC | 1,09 | 0,14 | EE vs. EC | 0,011 | -3,08 | |
| *Esr1* | | | CC | 1,00 | 0,20 | CC vs. CE | 0,001 | 3,09 | 7d |
| | | | CE | 0,29 | 0,26 | CC vs. EE | 0,002 | 3,15 | |
| | | | EE | 0,32 | 0,23 | CE vs. EC | 0,002 | -2,15 | |
| | | | EC | 0,97 | 0,36 | EE vs. EC | 0,003 | -2,12 | |
| *Oxt* | | | CC | 1,00 | 0,20 | CE vs. EC | 0,003 | 1,49 | 7e |
| | | | CE | 0,57 | 0,11 | EE vs. EC | 0,003 | 2,63 | |
| | | | EE | 0,56 | 0,16 | | | | |
| | | | EC | 1,27 | 0,52 | | | | |
| *Th* | | | CC | 1,00 | 0,25 | CC vs. EE | 0,000 | 3,22 | 7f |
| | | | CE | 1,00 | 0,22 | CC vs. EC | 0,000 | 2,89 | |
| | | | EE | 0,37 | 0,12 | CE vs. EE | 0,000 | 3,50 | |
| | | | EC | 0,32 | 0,22 | CE vs. EC | 0,000 | 3,05 | |
| *Kiss1* p H3K27me3 | | | CC | 1,00 | 0,18 | CC vs. EE | 0,004 | -2,69 | 7c |
| | | | CE | 2,18 | 0,79 | CC vs. CE | 0,011 | -2,05 | |
| | | | EE | 2,33 | 0,67 | EE vs. EC | 0,005 | -2,17 | |
| | | | EC | 1,04 | 0,50 | CE vs. EC | 0,015 | 1,72 | |
| *Esr1* p H3K9ac | | | CC | 1,00 | 0,41 | CC vs. EE | 0,046 | 1,81 | 7d |
| | | | CE | 0,41 | 0,17 | CC vs. CE | 0,039 | 1,86 | |
| | | | EE | 0,43 | 0,17 | | | | |
| | | | EC | 0,94 | 0,51 | | | | |
| *Oxt* p H3K4me3 | | | CC | 1,00 | 0,48 | CC vs. EE | 0,044 | 1,54 | 7e |
| | | | CE | 0,55 | 0,27 | EE vs. EC | 0,003 | 2,78 | |
| | | | EE | 0,44 | 0,20 | CE vs. EC | 0,012 | -2,19 | |
| | | | EC | 1,24 | 0,35 | | | | |
| *Th* p H3K27me3 | | | CC | 1,00 | 0,24 | CC vs. EE | 0,000 | -4,46 | 7f |
| | | | CE | 0,96 | 0,23 | CC vs. CE | 0,000 | 0,17 | |
| | | | EE | 2,11 | 0,26 | EE vs. EC | 0,000 | -0,12 | |
| | | | EC | 2,07 | 0,48 | CE vs. EC | 0,000 | -2,91 | |

**Notes**: Effect size was calculated using Cohen's *d*. F= generation. N=sample size. SD: standard deviation. IP: interpulse interval.
